# Feedback scales the spatial tuning of cortical responses during both visual working memory and long-term memory

**DOI:** 10.1101/2024.04.11.589111

**Authors:** Robert Woodry, Clayton E. Curtis, Jonathan Winawer

**Affiliations:** Department of Psychology, New York University, New York City, NY 10003; Center for Neural Science, New York University, New York City, NY 10003

**Keywords:** working memory, long-term memory, fMRI, retinotopy, saccades

## Abstract

Perception, working memory, and long-term memory each evoke neural responses in visual cortex. While previous neuroimaging research on the role of visual cortex in memory has largely emphasized similarities between perception and memory, we hypothesized that responses in visual cortex would differ depending on the origins of the inputs. Using fMRI, we quantified spatial tuning in visual cortex while participants (both sexes) viewed, maintained in working memory, or retrieved from long-term memory a peripheral target. In each condition, BOLD responses were spatially tuned and aligned with the target’s polar angle in all measured visual field maps including V1. As expected given the increasing sizes of receptive fields, polar angle tuning during perception increased in width up the visual hierarchy from V1 to V2, V3, hV4, and beyond. In stark contrast, the tuned responses were broad across the visual hierarchy during long-term memory (replicating a prior result) and during working memory. This pattern is consistent with the idea that mnemonic responses in V1 stem from top-down sources, even when the stimulus was recently viewed and is held in working memory. Moreover, in long-term memory, trial-to-trial biases in these tuned responses (clockwise or counterclockwise of target), predicted matched biases in memory, suggesting that the reinstated cortical responses influence memory guided behavior. We conclude that feedback widens spatial tuning in visual cortex during memory, where earlier visual maps inherit broader tuning from later maps thereby impacting the precision of memory.

**Significance Statement:** We demonstrate that remembering a visual stimulus evokes responses in visual cortex that differ in spatial extent compared to seeing the same stimulus. Perception evokes tuned responses in early visual areas that increase in size up the visual hierarchy. Prior work showed that feedback inputs associated with long-term memory originate from later visual areas with larger receptive fields resulting in uniformly wide spatial tuning even in primary visual cortex. We replicate these results and show that the same pattern holds when maintaining in working memory a recently viewed stimulus. That trial-to-trial difficulty is reflected in the accuracy and precision of these representations suggests that visual cortex is flexibly used for processing visuospatial information, regardless of where that information originates.

## Introduction

While visual maps in occipital cortex have been known to be essential for vision for over a century (Inouye, 1909), recent evidence supports the provocative idea that these visual maps also play important roles in both working and long-term memory. Surprisingly, the contents of visual working memory (Curtis & Sprague, 2021; Harrison & Tong, 2009; Serences et al., 2009) and long-term memory (Bosch et al., 2014; Naselaris et al., 2015; Vo et al., 2022) can be decoded from the patterns of voxel activity in human primary visual cortex (V1). Such results support influential theories of how the encoding mechanisms used for perception might also be used to store working memory representations (D’Esposito & Postle, 2015; Serences, 2016) and recall visual properties from long-term memory (Rugg et al., 2008; Schacter et al., 1998; Tulving & Thomson, 1973). It is no longer a question as to *whether* visual cortex participates in cognitive functions beyond perception, but a question of *how*.

Patterns of evoked activity during perception can be used to predict the contents of memory (Albers et al., 2013; Bosch et al., 2014; Rademaker et al., 2019), supporting the idea that stimulus features are encoded in visual cortex activity in a similar manner in perception and memory. Moreover, recalling large objects evokes activity that encroaches into more peripheral portions of visual field maps (Kosslyn et al., 1995), as if recalled objects that are larger encompass more of the visual field just like it does for seen objects. Despite these parallels, we hypothesized that these responses in striate and extrastriate cortex also differ because the origins of their inputs differ. During perception, visual information is transmitted through the eyes and the retinogeniculate pathway to a cluster of retinotopically organized maps, including V1, V2, and V3 (Van Essen & Maunsell, 1983). Neural activity in these early visual maps strongly influences one’s perception (Tong et al., 1998). During working memory, information enters visual cortex in the same feedforward way, but after stimulus offset the maintenance of visual information depends on interactions between higher order cortical regions like the prefrontal cortex and sensory areas like V1. Such interactions support the storage of working memory representations (Curtis & D’Esposito, 2003; Curtis & Sprague, 2021; D’Esposito & Postle, 2015). During long-term memory, information is thought to be reconstructed in visual cortex through the retrieval of a memory stored in other brain structures such as the hippocampus (Schacter et al., 1998).

Most of the research described above focused on similarities between memory and perceptual representations. Some recent work, however, has observed systematic differences in visual cortex activity between perception and long-term memory (Breedlove et al., 2020; Favila et al., 2022), and between perception and working memory (Chunharas et al., 2023; Duan & Curtis, 2024; Kwak & Curtis, 2022). Several important questions remain. There have been no comparisons of visual cortex responses in perception, working memory, and long-term memory with the same study parameters, no direct estimates of tuning width in visual cortex during working memory, and limited links between the precision of long-term memory reactivation and single-trial behavior.

Here, we use the receptive field properties of neural populations within voxels (Dumoulin & Wandell, 2008) in visual field maps to quantify and compare the spatial tuning of responses during perception and memory. As a preview, fMRI responses during perception matched the spatial position of seen targets and increased in tuning width up the visual hierarchy consistent with increases in receptive field sizes (Dumoulin & Wandell, 2008; Smith et al., 2001). During working and long-term memory, tuning widths were large in early visual cortex as if they were inherited from feedback from higher order areas with large receptive fields. This confirms studies of long-term memory (Breedlove et al., 2020; Favila et al., 2022) and shows now that the same pattern holds in working memory. Critically, errors in spatially tuned responses during visual memory aligned with errors in memory behavior, suggesting that memory behavior depends on these responses in early visual cortex.

## Materials and Methods

### Subjects

Eight human subjects (5 Males, 25-32 years old) were recruited to participate in the experiment and were compensated for their time. Subjects were recruited from the New York University community and included author R.F.W. Other subjects were naive to the purpose of the experiments. All subjects gave written informed consent to procedures approved by the New York University Institutional Review Board prior to participation. All subjects had normal or corrected-to-normal visual acuity, normal color vision, and no MRI contraindications. No subjects were excluded from the main data analyses.

### Stimuli

#### Target stimuli

For each of the three conditions (perception, working memory, long-term memory), 16 polar angles were selected from 16 evenly spaced bins along an isoeccentric ring at 7° eccentricity from a central fixation point (0-22.5 deg, 22.5-45 deg, … 337.5 to 360 deg). Within each bin, the precise target location was randomly assigned. The 48 target locations were uniquely generated for each participant. The target stimulus at each spatial location consisted of a drifting Gabor patch (sd = 0.33 deg, spatial frequency: 2 cyc/deg, truncated at 5 sd). Each Gabor patch drifted radially towards fixation at a rate of 3 Hz (1.5 deg/s), with an orientation tangential to the isoeccentric ring.

#### Object stimuli

A small image of an object (1° diameter) was shown at fixation at the beginning of each trial. For the long-term memory condition, each object stimulus paired with one of the 16 target locations. These object-target pairings remained consistent throughout the pre-scan behavioral training (see below) and during the main experiment, and therefore a particular object served as a cue for the location to retrieve from long-term memory. For the perception and working memory conditions, the object-target pairings were unique for each trial, with each object only ever presented once for each subject, ensuring that subjects could not form associations between objects and target locations. Because the long-term memory condition required pairing target Gabors to fixation stimuli that were easy to learn, and because the perception and working memory conditions required the presentation of a unique fixation stimulus on every trial, this required a large number of easily recognizable cues with a low probability of confusion. Therefore the fixation stimuli were selected randomly without replacement from a bank of colored images consisting of everyday objects (BOSS dataset, (Brodeur et al., 2010). For each subject a total of 144 unique everyday objects were sampled without replacement from this dataset, 16 were assigned to the long-term memory condition, and the other 128 assigned to each trial in the perception and working memory conditions.

### Experimental procedure

#### Main experiment

The main experiment cycled through three scans: one each for perception, working memory, and long-term memory, repeated twice per scan session (Figure 1). Each subject participated in two scan sessions spaced no more than a week apart, for a total of 12 scans (4 per condition). Each scan had 16 trials corresponding to the 16 target locations, in random order. Across both sessions, this culminated in a total of 64 trials per condition, 192 trials total per subject. The subjects were unaware of the trial design.

**Figure 1.**
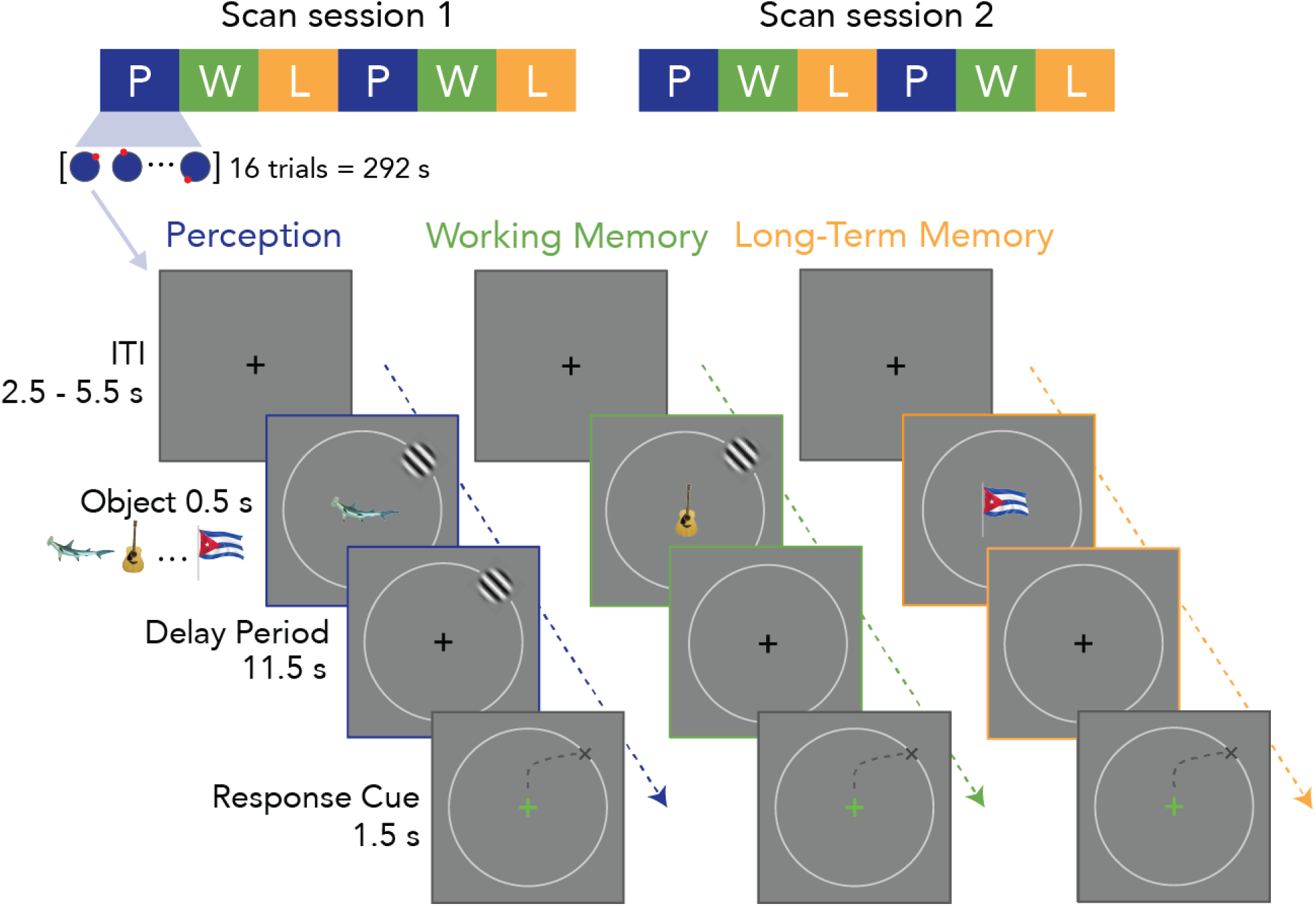
Main fMRI task design. Subjects participated in two fMRI sessions with 6 scans each: perception, working memory, and long-term-memory scans, repeated twice (top). Each 292-s scan included 16 trials corresponding to 16 polar angles, in random order, with the Gabor targets always at 7° eccentricity. Objects are shown centrally for 0.5 seconds, but are only meaningful for the long-term memory trials, for which they serve as cues for the target. In Perception trials, the Gabor target remains on screen for the 11.5-s delay. For working memory trials, the Gabor target disappears after 0.5 s. For long-term memory trials, the Gabor target is not shown at all. At the end of the delay, the fixation cross turns green and participants make a saccade to the location of the Gabor target. The conditions are matched except for how the Gabor target location is accessed and how it is maintained throughout the delay period.

For perception trials, we instructed participants to fixate on a central cross. The cross was briefly replaced by an object for 0.5 seconds. After the object disappeared, the cross re-appeared and the target stimulus remained visible for 11.5 seconds, during which subjects maintained central fixation. After this delay period, the fixation cross changed from black to green, indicating to the subject to make a saccade to the target location. Participants were instructed to maintain their fixation at the expected location until the fixation cross changed color back to black, at which point they returned their gaze to the central fixation cross. This saccade response period lasted 1.5 seconds for each trial.

The working memory block was the same except the target stimulus disappeared when the object disappeared. The subject was instructed to “hold the target in mind” throughout the delay while centrally fixating. At the end of the delay, the subject was similarly cued to make a saccade response to the location of the target stimulus.

The long-term memory block was the same except that the target stimulus was not shown at all. The subject was instructed to retrieve from memory the target stimulus associated with the object, with these associations learned during a pre-scan behavioral training session (see next section). Similar to the working memory and perception blocks, participants maintained central fixation during the delay period of 11.5 seconds and made a saccade to the target location when the cross turned green.

#### Long-term memory training

Subjects learned associations between 16 object stimuli and 16 target locations by completing study and retrieval blocks before each scan session (outside the scanner). Following each study block, subjects completed three retrieval blocks. In each trial of the study block, an object stimulus was briefly presented at fixation simultaneously with its corresponding Gabor target stimulus (Figure 2a). Subjects were instructed to fixate a central cross and learn the association between each object stimulus and its corresponding target location. The study block was self-paced with a minimum 1 second inter-trial interval (ITI). Each of the 16 object/target pairs was presented five times per block (80 trials), with at least two study blocks per behavioral training session.

**Figure 2.**
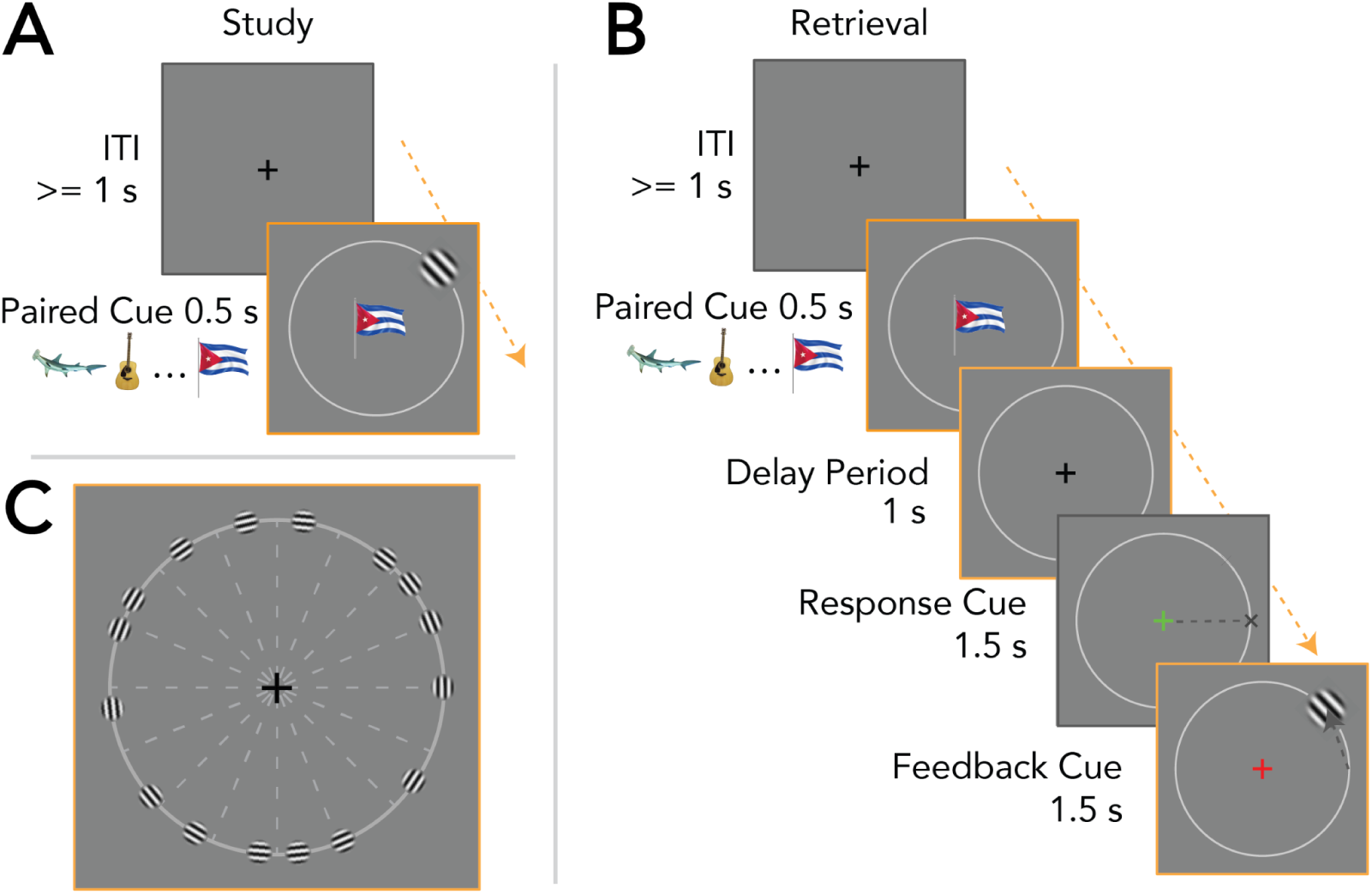
Prescan long-term memory training. A) Study phase: Self-paced brief (0.5 second) viewings of the paired pre-cue at fixation and the associated target Gabor in the periphery, followed by an inter-trial interval (min. one second) Each participant had their own set of 16 unique pairs to memorize. B) Retrieval phase: The bulk of learning happened during this training phase. The retrieval phase was similarly self-paced, where following an inter-trial interval (min. one second), a paired cue was briefly presented at fixation without the target in the periphery. After a brief delay period the participant was cued to make a saccade response to the target’s location. Feedback was then given in the form of i) the fixation cross either remaining green if the saccade was closest to the target’s location, or changing to red if the response landed closer to a different target’s location, and ii) a brief presentation of the peripheral target Gabor, irrespective of accuracy. C) Example layout of all target stimuli shown in a single image: For each condition, target stimuli are presented at 7° eccentricity, sampled around the visual field within each of 16 non-overlapping 22.5° bins. Every participant had their own unique set of target locations. This led to some targets spaced near each other (i.e. near the bin borders marked by the dashed lines), while others were spaced far apart. Note that only a single target was shown in each trial, as in Panel B.

In the retrieval block, subjects were instructed to fixate the central cross while the object stimulus was flashed briefly, followed by a short delay where the subject was instructed to retrieve from memory the associated target stimulus (Figure 2b). This was followed by the fixation cross changing color to green, which indicated to the participant to make an eye movement to the target’s expected location. Feedback followed each retrieval trial in the form of the fixation cross remaining green if correct or changing color to red if incorrect. A saccade was considered correct if it was closer to the correct target than to any of the other 15 possible targets, otherwise it was considered incorrect. Simultaneously, the target stimulus was revealed at its true location, with the subject instructed to make a corrective saccade to the target stimulus. The retrieval block was also self-paced with a 1 second minimum ITI. Each of the 16 object/target pairs was presented four times per block (64 trials), with at least six retrieval blocks for the first training session, and four for the second session. Subjects were asked after six blocks if they felt they learned all the associations or if they needed to do more practice. If they needed more practice, they did at least one more retrieval block.

#### Retinotopic mapping procedure

In addition to the main experiment each participant completed 10-12 retinotopic mapping scans in a single separate scan session. The stimuli and procedures are the same used by Himmelberg et al. (2021), described here in brief. Each scan consisted of contrast patterns windowed by bar apertures (1.5 deg width) that swept across the visual field within a 12-deg-radius circle. There were 8 sweeps along different directions, 4 cardinal (left, right, up, down), and 4 diagonal. The cardinal sweeps traveled the entire extent of the circular aperture in 24 s. The diagonal sweeps stopped halfway (12-s), and were then replaced by blank periods of 12-s. Hence each bar sweep took 24 s to complete. At 8 sweeps for each functional run, each scan took 192 s in total to complete. The contrast patterns were pink noise (grayscale) backgrounds with randomly placed and sized items. The stimuli and background were updated at 3 Hz. Participants were instructed to report any observed change in fixation dot color with a button box press. Color changes occurred around once every three seconds. The contrast patterns for the mapping stimuli were first used by Benson et al (2018).

### MRI acquisition

Imaging was conducted at the Center for Brain Imaging at New York University using a 3T Siemens Prisma MRI system and a Siemens 64-channel head/neck coil. We acquired functional images with a T2*-weighted multiband echo planar imaging (EPI) sequence with whole-brain coverage (repetition time = 1 s, echo time = 37 ms, flip angle = 68°, 66 slices, 2 x 2 x 2 mm voxels, multiband acceleration factor = 6, phase-encoding = posterior-anterior). We collected spin echo images with anterior-posterior and posterior-anterior phase-encoding to estimate, and correct for, the susceptibility-induced distortion in the functional EPIs. We also acquired one to three whole-brain T1-weighted MPRAGE 3D anatomical volumes (.8 x .8 x .8 mm voxels) for each of the eight subjects.

### MRI processing

All original MRI data (DICOM files) were defaced to anonymize them using *pydeface* (https://github.com/poldracklab/pydeface). The DICOM data were then converted to NIFTI and organized into the Brain Imaging Data Structure format (K. J. Gorgolewski et al., 2016) using Heuristic Dicom Converter (Halchenko et al., 2018). The data were then preprocessed using fMRIPrep 20.2.7 (Esteban et al., 2018, 2019), which is based on Nipype 1.7.0 (K. Gorgolewski et al., 2011; K. J. Gorgolewski et al., 2018).

#### Anatomical data preprocessing

The following sections on anatomical and functional data preprocessing are provided by the fMRIPrep boilerplate text generated by the preprocessed scan output.

Each of the one to three T1w images was corrected for intensity non-uniformity with N4BiasFieldCorrection (Tustison et al., 2010), distributed with ANTs 2.3.3 (Avants et al., 2008). The T1w-reference was then skull-stripped with a Nipype implementation of the antsBrainExtraction.sh workflow (from ANTs), using OASIS30ANTs as target template. Brain tissue segmentation of cerebrospinal fluid, white-matter and gray-matter was performed on the brain-extracted T1w using fast (FSL 5.0.9, (Zhang et al., 2001)). A T1w-reference map was computed after registration of the T1w images (after intensity non-uniformity-correction) using mri_robust_template (FreeSurfer 6.0.1, (Reuter et al., 2010)). Brain surfaces were reconstructed using recon-all (FreeSurfer 6.0.1, (Dale et al., 1999)), and the brain mask estimated previously was refined with a custom variation of the method to reconcile ANTs-derived and FreeSurfer-derived segmentations of the cortical gray-matter of Mindboggle (Klein, 2017).

#### Functional data preprocessing

For each of the 12 BOLD runs found per subject (across all tasks and sessions), the following preprocessing was performed. First, a reference volume and its skull-stripped version were generated by aligning and averaging a single-band reference. A B0-nonuniformity map (or fieldmap) was estimated based on two EPI references with opposing phase-encoding directions, with 3dQwarp (Cox & Hyde, 1997). Based on the estimated susceptibility distortion, a corrected EPI reference was calculated for a more accurate co-registration with the anatomical reference. The BOLD reference was then co-registered to the T1w reference using bbregister (FreeSurfer) which implements boundary-based registration (Greve & Fischl, 2009). Co-registration was configured with six degrees of freedom. Head-motion parameters with respect to the BOLD reference (transformation matrices, and six corresponding rotation and translation parameters) are estimated before any spatiotemporal filtering using mcflirt (FSL 5.0.9, (Jenkinson et al., 2002)). BOLD runs were slice-time corrected to 0.445s (0.5 of slice acquisition range 0s-0.89s) using 3dTshift from AFNI 20160207 (Cox & Hyde, 1997). First, a reference volume and its skull-stripped version were generated using a custom methodology of fMRIPrep. The BOLD time-series were resampled onto the fsnative surface. The BOLD time-series (including slice-timing correction) were resampled onto their original, native space by applying a single, composite transform to correct for head-motion and susceptibility distortions. These resampled BOLD time-series will be referred to as preprocessed BOLD. All resamplings can be performed with a single interpolation step by composing all the pertinent transformations (i.e. head-motion transform matrices, susceptibility distortion correction, and co-registrations to anatomical and output spaces). Gridded (volumetric) resamplings were performed using antsApplyTransforms (ANTs), configured with Lanczos interpolation to minimize the smoothing effects of other kernels (Lanczos, 1964). Non-gridded (surface) resamplings were performed using mri_vol2surf (FreeSurfer).

Many internal operations of fMRIPrep use Nilearn 0.6.2 (Abraham et al., 2014), mostly within the functional processing workflow. For more details of the pipeline, see the section corresponding to workflows in fMRIPrep’s documentation.

#### GLM analyses

From each subject’s surface based time series, we used GLMSingle (Prince et al., 2022) to estimate the neural pattern of activity during the 11.5-s delay periods of the main experiment for each trial. GLMSingle is a three step process where 1) an optimal hemodynamic response function (HRF) is fit to each vertex’s time series from a bank of 20 HRF functions obtained from the Natural Scenes Dataset (Allen et al., 2021) via an iterative linear fitting procedure. 2) Noise regressors are computed from the data by identifying noisy vertices defined by negative *R^2^,* deriving noise regressors from this noise pool using principal component analysis, then iteratively removing each noise regressor from all vertices’ time series. The optimal number of regressors is determined via a cross-validated *R^2^* improvement for the task-model. 3) GLMSingle implements fractional ridge regression as a way to improve robustness of single-trial beta estimates, particularly useful here as our design yields a limited number of trials per target position within each condition.

We constructed our design matrices to have 48 regressors of interest (16 polar angle bins x 3 conditions), with the events modeled as boxcars corresponding to the 11.5 s delay periods. We estimated one model for each participant with these designs in GLMSingle, resulting in single trial estimates for each trial in each surface vertex.

#### Fitting pRF models

Using the data from the retinotopy session, we fit population receptive field (pRF) models for each vertex on the cortical surface, as described by Himmelberg et al. (2021; section 2.6). In brief, for each surface vertex we fit a circular 2D-Gaussian linear population receptive field (pRF) to the BOLD time series, averaged across identical runs of the bar stimulus. The software was implemented in *Vistasoft* as described in Dumoulin & Wandell (2008), with a wrapper function to handle surface data (https://github.com/WinawerLab/prfVista). The models are parameterized by the Gaussian center (*x, y*) and standard deviation (σ).

### Visual field map definitions

Visual field maps were defined by drawing boundaries at polar angle reversals on each subject’s cortical surface using an early version of the visualization tool, cortex-annotate (https://github.com/noahbenson/cortex-annotate), which is built on *neuropythy* software (https://github.com/noahbenson/neuropythy, (Benson & Winawer, 2018). We followed common heuristics to define seven maps spanning early to mid-level visual cortex: V1, V2, V3 (Benson et al., 2022; Himmelberg et al., 2021); hV4 (Winawer & Witthoft, 2015); V3A and V3B (grouped into one ROI, V3ab) and IPS0 (Mackey et al., 2017); and LO1 (Larsson & Heeger, 2006).

We defined experiment-specific regions of interest for each visual field map composed of vertices whose pRF centers were near the target eccentricity and whose variance explained by the pRF model was above 10%. Specifically, we included only those vertices whose pRF centers were within one σ of 7° (the target eccentricity in the experiments). For example, a vertex with pRF center at 6 deg and pRF size (σ) of 1.5 deg would be included, but a vertex with pRF center at 6 deg and pRF size of 0.5 deg would not be included. We imposed the eccentricity restriction for the purpose of examining polar angle activation profiles, described in the next section. These measures are based on the retinotopy scans only and are therefore independent of the main experiment.

### Analyses quantifying perception, working memory, and long-term memory activity

To examine the BOLD response within the 11.5 second delay, we constructed polar-angle activation profiles for each visual field map and each condition (perception, working memory, long-term memory). Some analyses averaged over the delay period. For these analyses, we obtained the response amplitudes from GLMsingle (one beta weight per trial for each surface vertex). The visual field coordinates for each vertex came from pRF mapping, which was conducted in a separate fMRI session. We binned the response amplitudes by the polar angle distance between each vertex’s pRF and the target location on that trial. Binning by polar angle distance from the target enabled us to average across trials with different target locations, resulting in an activation profile as a function of distance from the target. This results in a distinct polar angle activation profile for each subject, condition, and visual field map. Prior to averaging across subjects, we normalized each activation profile by dividing by the vector length of the perceptually evoked activation profile for that map. To preserve meaningful units, we then rescaled the activation profile by the average of vector lengths across subjects. Visual inspection of the average profiles showed a peak near 0, and negative responses far from 0. We fit the functions with a difference of two von Mises distributions, constrained so that the centers were the same.

A separate procedure was used to derive 2D activation profiles, which include time as well as polar angle. To derive these, we extracted the denoised BOLD time series for each vertex on each trial, rather than a single beta weight per trial, expressed as percent signal change from the mean of each scan. The denoised data were derived from the output of a GLM analysis using the GLMdenoise algorithm (Kay, Rokem, et al., 2013). This algorithm is closely related to the GLMSingle algorithm used for the static analyses, but has greater flexibility in specifying the trial structure (e.g., events with different durations) than the GLMSingle algorithm, but doesn’t return estimates for single trials. The event beta weights from this GLM were not used for any analyses; instead we only used it to derive denoised BOLD time series for subsequent temporal analyses.

For these 2D analyses, time was sampled at seconds from -3 s to 17 s in each trial, relative to the start of the delay period. We then computed polar angle activation profiles independently for each time point, using the procedure described above: binning by polar angle distance from the target, averaging across trials within a condition and visual field map. This results in 2D activation profiles, which span polar angle and time within a trial.

For temporal analyses these 1D polar angle activation profiles were computed at each timepoint (TR = 1 sec) from the three seconds before the onset of the paired cue to 17 seconds after, resulting in a 2D heatmap of the polar angle activation profiles over time. As in the static activation profiles, the vertices were binned by polar angle distance from the target, and averaged across trials within each visual map, separately for each condition and subject. This results in a matrix that is polar angle by time. We fit the polar angle profile independently for each time point, again using a difference of Von Mises’ distributions, constrained so that the centers were the same.

For both analyses, we estimated the amplitude (trough to peak), the peak location, and full-width-at-half-maximum (FWHM) from each of the difference of Von Mises fits. We bootstrapped across subjects (with replacement) 10,000 times to obtain 68% and 95% confidence intervals for these location, amplitude, and FWHM parameter estimates.

#### Temporal analyses

We characterized the time course for each 2D polar angle activation profile by fitting logistic functions to the amplitude estimate at each time point from -3 seconds to 14 seconds relative to the onset of the paired cue. This is because, after 14 seconds, the saccade response dominated the signal introducing noise to the target-aligned response. The logistic fits resulted in estimates of four parameters for each 2D polar angle activation profile: *t_0_*, the rise-time to reach the function’s midpoint; *L,* the upper asymptote of the amplitude estimates; *k*, the logistic growth rate of the function (i.e. the steepness of the curve); and *c* a baseline. All four parameters were constrained to have a lower bound of 0; the upper bound was unconstrained for *L* and *c,* 15 seconds for *t_0_*, and 5% signal change / second for *k*. This fit was sufficient for the perception and long-term memory conditions, which generally showed a rise and then a steady response. For working memory, the response rose transiently, and then declined to a lower value. This pattern was accurately captured by fitting a multiplication of two logistic functions rather than a single logistic function. The two functions were constrained to have the same *L* parameter, but could differ in *t_0_, c*, and *k*. The parameter bounds for the second logistic function were the same as the first, except where *k* was constrained to be negative instead of positive. We repeated these logistic fits for each of the 10,000 bootstraps, computing 68% confidence intervals of the estimated logistic time series.

#### Saccade analyses

We used 2 EyeLink eye trackers (SR Research, Ottawa, ON, Canada) with a 1000-Hz sampling rate, one in the scanner and one outside the scanner. In the scanner the EyeLink 1000 plus was mounted onto a rig in the magnet bore that holds the projection screen. In the psychophysics room for long-term memory training, we used an EyeLink Tower mount with a 25-mm lens mounted on the EyeLink camera to allow close viewing.

During both the training and the fMRI experiment, saccades were labeled by the EyeLink’s default saccade detector, which classifies saccades as eye movements with velocity and acceleration exceeding 30 deg/s and 8000 deg/s^2^, respectively. Saccade responses were collected during the response window at the end of each trial. For all behavioral analyses, the saccade responses used are those which landed nearest the target eccentricity during the saccade response window.

Subjects often make multiple saccades to get to the target. We defined the endpoint as the saccade whose eccentricity was closest to the target eccentricity (7°), irrespective of the polar angle. We then measured the angular distance between this point and the target (ignoring eccentricity). We excluded saccade responses whose eccentricity was less than 3.5° or greater than 12° visual angle from fixation. One subject was removed from the saccade analysis due to technical error with the eye tracker during the scan sessions. Data from three scans from another subject were also excluded due to calibration error.

For comparison between BOLD data and saccades, we divided the saccade data for each subject and each condition into tertiles for counterclockwise, center, and clockwise. We repeated the ROI-level analyses on this split saccade data to obtain 1D polar angle activation profiles for each ROI, condition, subject, and tertile.

#### Resampling Statistics

We used the bootstrapped data from each analyses to make inferences on spatial tuning properties as a function of condition and of other trial-level factors. Statistics reported are computed using the bootstrapped data. To assess our main claims, we report the mean and confidence intervals from the bootstrapped data. For comparisons between a measurement and a fixed value, we report the 95% CI, which corresponds to ±2 standard deviations of a normal distribution. For comparisons between two estimates, we report the 68% CIs, which corresponds to ±1 standard deviation of a normal distribution.

To assess the relationship between a map’s position in the visual hierarchy and spatial tuning width, we assigned each map an ordinal value according to its relative position: 1, V1; 2, V2; 3, V3; 4, hV4, LO1, V3A/B; 5, IPS0. We fit a line of tuning width vs ordinal position for each bootstrap, generating a distribution of slope means for each condition. The rest of our analyses took the form of computing the differences of mean effects between conditions, both for individual visual maps and for early (V1-V3) vs. later (hV4-IPS0) visual cortex.

### Software

Data visualization, model fitting, and statistical quantification for all analyses described in this paper were made using matplotlib 3.5.2 (Hunter, 2007), nibabel 3.2.2 (Brett et al., 2022), pandas 1.4.2 (The pandas development team, 2024), scikit-learn 1.0.2 (Pedregosa et al., 2011), scipy 1.8.0 (Virtanen et al., 2020), and seaborn 0.11.2 (Waskom, 2021).

## Results

We tested how the spatial tuning of cortical visual representations is shaped by viewing a peripheral target, maintaining it in working memory, or retrieving it from long-term memory. We also tested how the cortical representations during memory relate to the metrics of memory-guided saccades.

### Responses in visual cortex are spatially tuned during working memory and long-term memory

We first ask whether activation in early visual cortex during memory is spatially tuned. To assess this, we parameterized the sensory representations generated during the delay period by remapping GLM estimates of brain activity from the cortical surface to visual space (Figure 3A,B). We then computed polar angle activation profiles to capture the spatial tuning of sensory representations generated during the delay period (Figure 3C), and estimated their amplitude, peak location, and tuning width (Figure 3D). We compared these estimates between perception, working memory, and long-term memory conditions.

**Figure 3.**
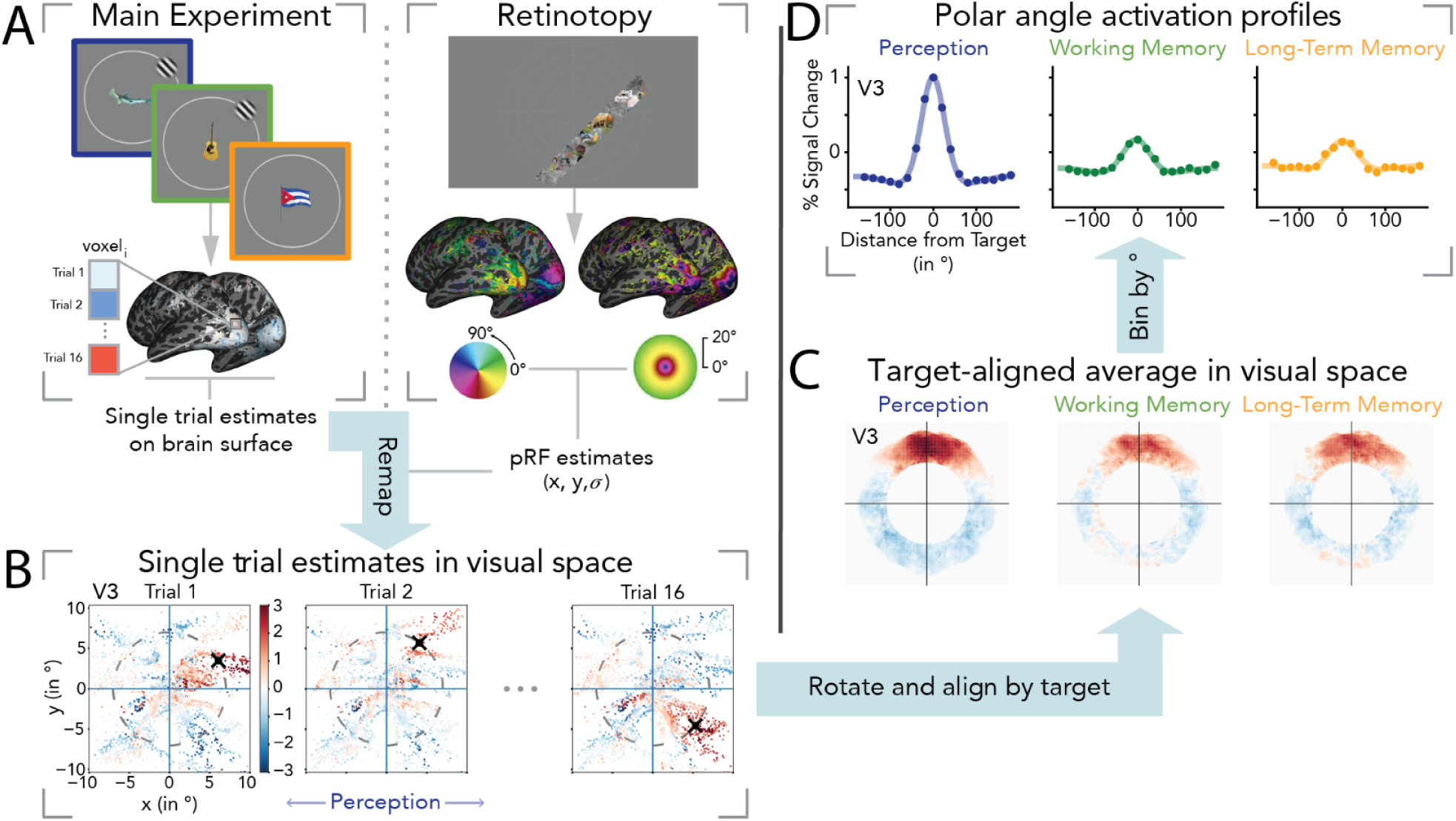
Target-aligned averages of sensory representations in visual space. (A) Single trial beta estimates for each vertex on the brain surface are obtained from the main task’s delay period. In addition, a separate retinotopic mapping procedure is used to obtain population receptive field (pRF) estimates for each vertex. (B) We use the pRF estimates to remap the single trial estimates on the brain surface to single trial estimates in visual space, that are then rotated and aligned by trial target position. The pRF estimates are also used to define retinotopic maps for seven regions of interest across visual cortex, and to restrict voxels to those with pRF centers near the target eccentricity. (C) Normalizing and averaging the aligned trial estimates for each map yields target-aligned averages in visual space for each condition. A 2D visualization of a subject’s target-aligned average is shown above for visual map V3. (D) Beta estimates are binned by polar angle distance from target and fit to a difference of Von Mises functions to produce polar angle activation profiles. This mapping of BOLD response as a function of polar angle distance from the stimulus captures the spatial tuning profile of cortical responses during perception, working memory, and long-term memory. Panels in this figure and all following figures can be reproduced using code contained in the /paper/figures folder at https://github.com/rfw256/Woodry_2024_Cortical-tuning-of-visual-memory/. The panels ‘Single trial estimates’ and ‘target-aligned averages’ are generated using *fig3_09-16-2024.py*.

The clearest difference between perception and the two memory conditions is that the BOLD amplitude is much larger during perception. This difference is particularly evident in earlier visual maps V1-V4. The larger response during perception was in part due to the drifting Gabor target, which further drove the response during the delay period. While the amplitudes were lower during memory, they were all positive, ranging from 0.25% to 0.5% percent signal change across visual maps (Figure 4B. middle). The 95% confidence interval did not overlap 0% BOLD response in any ROI.

**Figure 4.**
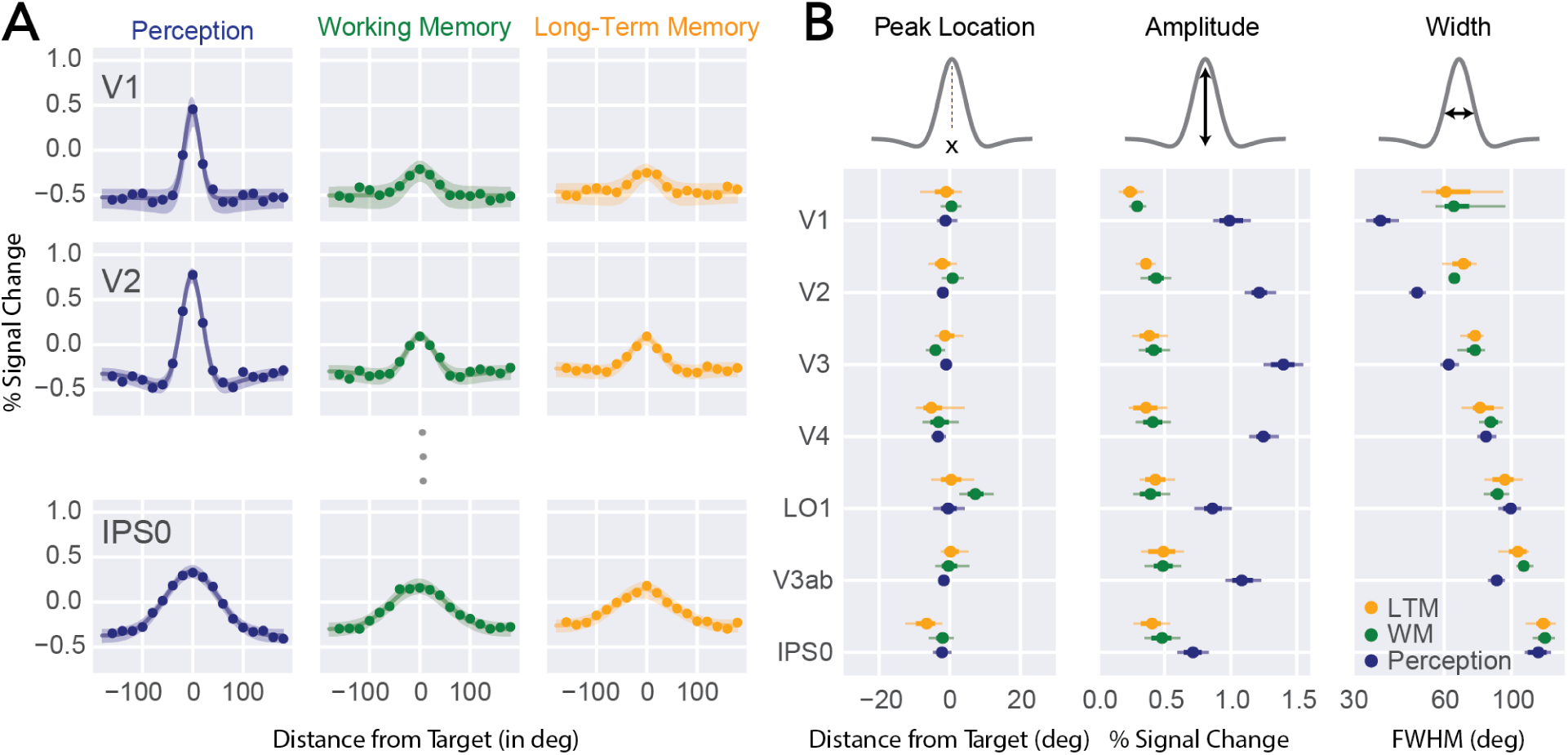
Memory has broader spatial tuning than perception in earlier visual cortex. A) Polar angle activation profiles fit to brain activity during 11.5 second presentation/delay period. Highlighted regions indicate 68% confidence intervals bootstrapped across subjects. Perception working memory, and long-term memory brain activity show spatial tuning to target locations across visual ROIs. B) Spatial tuning metrics obtained from polar angle activation profiles across visual ROIs. The dots are means across bootstraps. The thick shading is the 68% confidence interval. The thin shading is the 95% confidence interval. Dotted line in the leftmost panel represents the target location (0°). The panels here are generated using *fig4_09-16-2024.py*.

The memory activation profiles, like the perception activation profiles, were spatially tuned to the stimulus. Specifically, for all 3 conditions and all 7 visual field maps, peak location estimates were centered around 0° relative to the stimulus angle (Figure 4B, left). Polar angle activation profiles which peak at 0° indicate accurate tuning to the true target location. Tuning to the target location is of course expected in the perception condition. But remarkably, we even see tuning to the target location in the earliest visual map, V1, during both memory conditions. (The confidence intervals from all three conditions include 0°.) Tuning in the two memory conditions confirms prior work showing engagement of visual areas, including primary visual cortex, during memory (Breedlove et al., 2020; Favila et al., 2022; Sprague & Serences, 2013; Vo et al., 2022).

### Memory shows broader spatial tuning than perception in early visual cortex

The lower amplitude but similar peak location during memory compared to perception suggests that memory responses might be similar to perception except for a scale factor. This turns out to not be correct. We find that instead of memory responses resembling perception responses (up to a scale factor), the memory responses show a difference in tuning width, which cannot be achieved by simply increasing or decreasing the response amplitude.

During perception trials, tuning widths increased sharply from early to later visual maps. To quantify this tendency, we assigned each map an ordinal value based on its estimated position in the visual hierarchy: 1, V1; 2, V2; 3, V3; 4, hV4, LO1, V3A/B; 5, IPS0. We fit a line of tuning width vs ordinal position, and found that for perception, the tuning width increased about 21° per position in the hierarchy (slope = 20.9°, CI [18.5, 23.8]), consistent with what is known about receptive fields increasing in size from early to later visual areas (Smith et al., 2001); (Dumoulin & Wandell, 2008). In contrast, there was little increase in tuning width across visual maps during both forms of memory, with a slope nearly two-thirds that measured during perception (working memory: slope = 15.2°, CI [10.6, 18.2]; long-term memory: slope = 14.8°, CI [7.7 18.6]). Because the tuning widths were so similar for long-term memory and working memory, we compared the average of the two memory conditions to perception. This steeper slope for perception than memory was robust (diff = 6.2°, CI [3, 13.3]).

In early visual maps V1-V3, spatial tuning during memory was broader than during perception (diff = 20°, CI [15.8, 29.7]), especially in V1 (diff = 28.8°, CI [15.9, 61]; Figure 4B, right). In later maps – hV4, LO1, and IPS0 – tuning widths were about the same in perception, working memory, and long-term memory (diff = 0.31°, CI [-6.7, 8.5]). Therefore, the memory responses are not just a scaled down version of the perceptual responses, but rather show broader tuning in earlier maps.

Perhaps the apparent increased tuning width during memory comes from the choice of eccentricity band for the analysis. Breedlove et al. (2020) reported that a voxel’s preferred eccentricity for remembered stimuli is shifted foveally. Hence our analyses, which were limited to voxels near the target eccentricity of 7°, might have missed the largest responses during memory. To assess this possibility, we recomputed the polar angle response profiles assuming target eccentricities with lower or higher eccentricity. We found that over a broad range of eccentricities, especially further in the periphery, the pattern of results was the same. Specifically, from 7 deg (target eccentricity) to 12 deg (limit of our retinotopic mapping), the patterns were unchanged (Figure 5b). At lower eccentricities, the response amplitudes declined. Note that more foveal tuning during long-term memory, as suggested by Breedlove et al. (2020), would predict that the best response should be in voxels whose receptive fields are more peripheral than the stimulus. Hence these analyses confirm that the broader tuning in memory is robust to the exact eccentricity band used for analysis.

**Figure 5.**
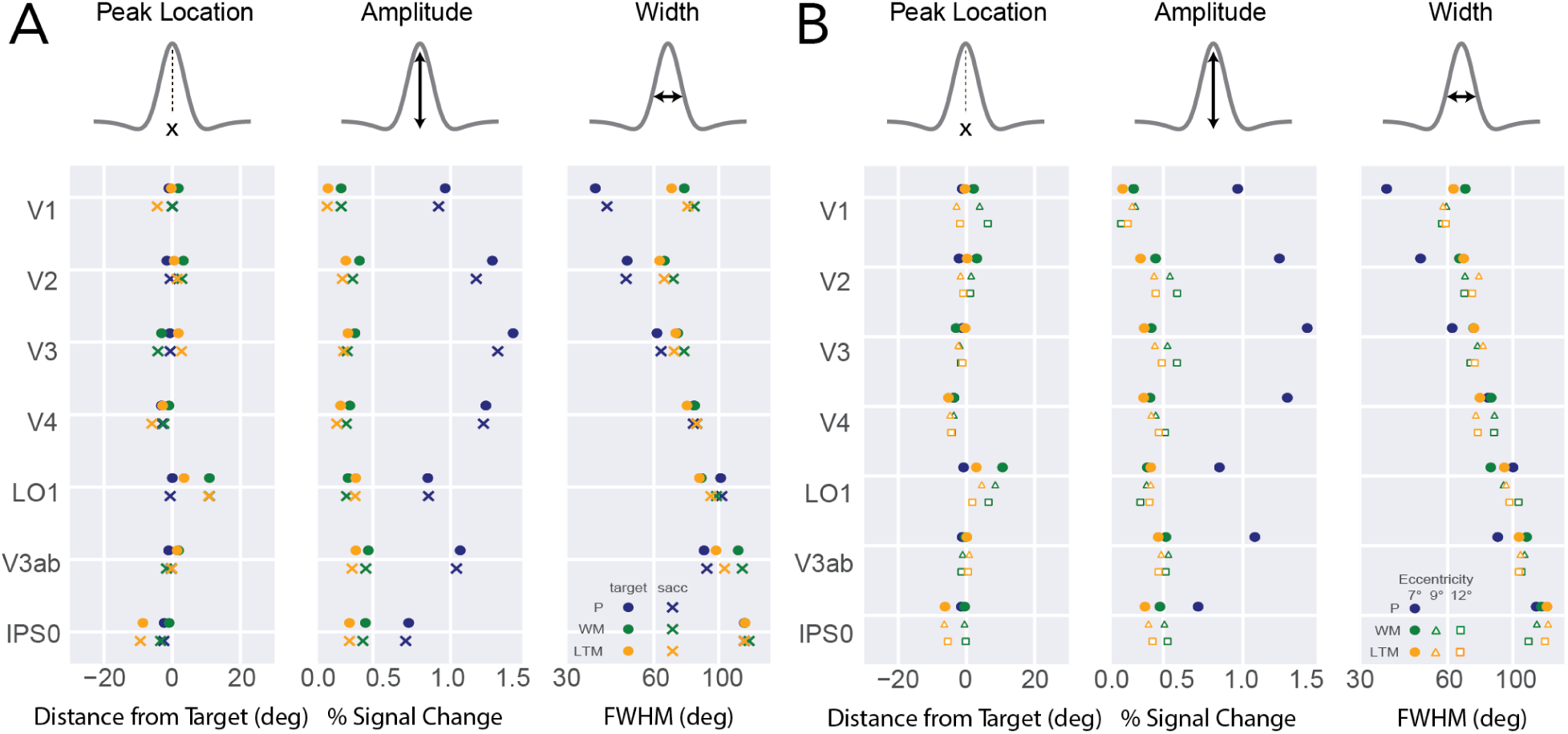
Broader spatial tuning during memory is robust when aligning to angular location of saccade responses or to different eccentricities. A) Tuning estimates as a function of assuming greater target eccentricities. While amplitude is higher at increased eccentricity bands, the broader tuning profiles during memory remain consistent. B) Aligning to the polar angle location of the saccade response produces spatial tuning estimates similar to the original analyses. This shows that the broader tuning observed when aligning the memory responses to the target are likely not due to error in memory recall. If so, this would predict a narrower memory response when aligning responses to the polar angle location of the saccade, assuming it is a reasonable proxy for the participants’ estimate of the target location.

A second possibility was that the broader tuning observed in early visual cortex is an artifact of averaging across trials: If tuning is as narrow during memory as in perception, but the tuning *center* varies from trial to trial more during memory than perception, then averaging data across trials would inflate the estimated width during memory. To assess this, we aligned each trial’s data to the polar angle of the saccade response rather than to the polar angle of the target, assuming that the saccade provides a reasonable estimate of where the participant thought the target was on each trial. Recomputing the tuning functions with this alignment did not alter the main pattern of results, again showing greater tuning width during memory than perception for early visual areas (Figure 5A). This argues against the possibility that the broader tuning in early visual cortex observed during memory is instead due to larger error during recall.

Because the only differences between conditions are how visual information is accessed, the observed differences in tuning widths confirm our hypothesis that differences in how information enters visual cortex shapes the spatial tuning of the subsequent representations.

### Distinct spatial tuning profiles emerge over time

The analysis above showed that the routing of stimulus information to visual cortex –feedforward in perception vs top-down in memory– affects the spatial representation. Here we ask how the routing affects the temporal dynamics of the response. Our expectation is that long-term memory responses will be slowest because the responses must be generated entirely internally (no stimulus is viewed). We can also ask whether working memory and long-term memory signals show the same tendency to sustain over the delay. To compare the temporal dynamics across conditions, we averaged the BOLD time series throughout the delay period across trials, binned by polar angle distance from the target (Figure 6a). We then computed the peak amplitude at each time point, and fit a rising logistic function (perception and long-term memory) or the product of a rising and falling logistic (working memory) across the time series (Figure 6b). The logistic product captures both the transient response evoked by the target at the beginning of the working memory delay period and the decay to a lower, sustained, activation for the remainder of the delay period. Because we fit the working memory response using a different function, we do not compare logistic parameters to the other two conditions. We make two observations that distinguish perception from long-term memory from working memory.

**Figure 6.**
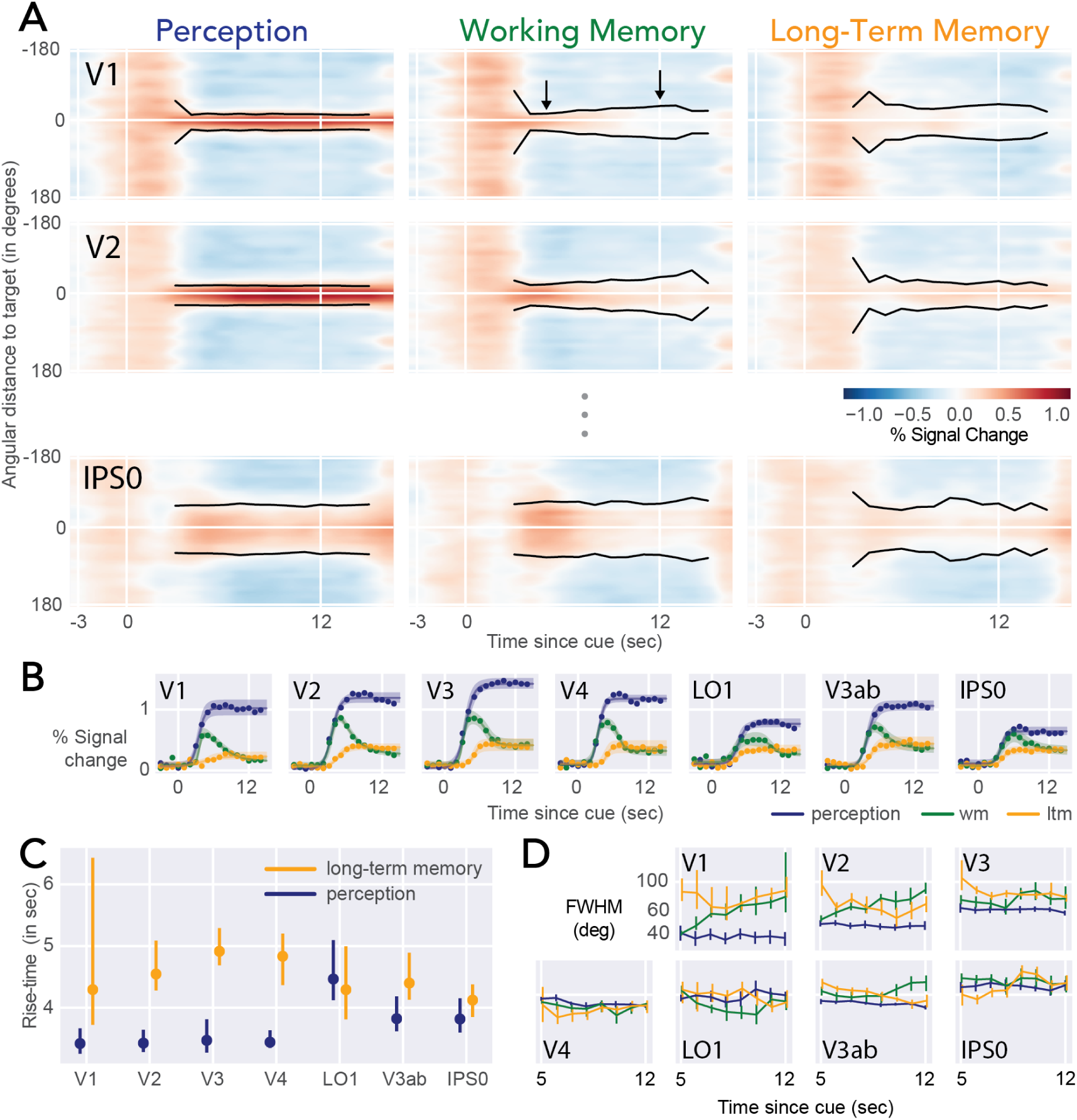
Perception, working memory, and long-term memory have distinct tuning over time. A) BOLD time course during the delay periods. Target location is 0°. Tuning width estimates are plotted as black lines, starting from 3 seconds after the onset of the delay. The arrows in the V1 working memory panel indicate the time at which the working memory amplitude peaks and the end of the delay period. The working memory responses in V1 and V2 are initially narrow, similar to perception (first black arrow), and then widen, similar to LTM (second black arrow). B) Logistic functions fitted to amplitude estimates over time. Shaded regions represent the 68% confidence interval of the logistic fits, bootstrapped across subjects. C) Comparison of rise-times to the inflection point of logistic fits between perception and long-term memory. Error bars represent 68% bootstrapped confidence intervals. D) Time course of tuning width (same values as black lines in panel a). Error bars are 68% bootstrapped confidence intervals. The panels here are generated using *fig5_09-16-2024.py*.

First, the working memory responses show a clear transition from a stimulus-driven transient, peaking at about 4 to 5 seconds after the cue, to a lower sustained signal. This is expected because the target stimulus is briefly shown prior to the delay during working memory. Corroborating this interpretation, the tuning width in early visual areas is narrow during working memory at 4 to 5 seconds after the cue, similar to the perceptual condition. It then increases in width by stimulus offset, so that the width at offset is similar to the width in long-term memory. The increase in width in V1 is 36.6° (CI [19°, 125.5°]) and in V2 is 35.3° (CI [28.2°, 45.4°]). This is indicated by the arrows in Figure 6A (for V1) and summarized in the time series in Figure 6D. Since the perceptually driven responses are brief, the estimates from the static analyses in Figure 4 are similar to those at the end of trial, visible in the dynamic plots of Figure 6.

In contrast to the working memory dynamics in V1 and V2, there is no clear change in the tuning width of the working memory response in other visual areas, except for in V3ab (diff = 18.1°, CI [4.3°, 28.3°]). This is to be expected given that the tuning widths of the working memory responses are matched to perception in these later visual areas. Moreover, there is little change in tuning width during the long-term memory trials or the perception trials.

Second, we find a general tendency for slower rise-time in long-term memory than perception across visual maps, with longer rise times in all maps except LO1, most prominently in earlier visual areas V1-V4 (diff = -1.4 sec, CI [-3.6, -0.3]; Figure 6c). This is consistent with memory responses arising later due to the sluggishness of feedback. Moreover, the rise times are much more variable in memory, as expected from responses that are internally and effortfully generated (memory) rather than from external, stimulus-triggered responses (perception).

### Errors in cortical tuning aligned with errors in memory-guided behavior

The results above demonstrate that visual cortex is engaged in a retinotopically specific way during working and long-term memory. These results do not, however, indicate whether these retinotopic representations are relevant for behavior. This is an important and open question in the field, with many reports claiming the representations are linked to behavior (Bone et al., 2019, 2020; Hallenbeck et al., 2021; Li et al., 2021) though some question their relevance (Xu, 2017). If the visual cortex representations are relevant for behavior, we expect alignment between the memory-driven cortical responses during the delay and subsequent saccade responses. Here, we took advantage of trial variability to test whether the peak location of cortical responses during the delay aligned with the direction of saccade error.

To test the alignment between cortical and saccade responses, for each subject we split the trials into three groups by their saccade error (Figure 7b), those trials with saccades near the targets, counterclockwise of the targets, or clockwise of the targets, with angular thresholds set to make the three bins contain equals number of trials. We then repeated our spatial tuning analyses separately for the clockwise and counterclockwise trials to compute the estimates of peak location during the delay period (Figure 7c). To reduce the number of comparisons, we defined a new region of interest as the union of all 7 maps. For this large ROI, there was a strong link between neural tuning and saccade error in long-term memory: the peak estimates for trials with clockwise saccades were 16.5° more clockwise than the trials with counterclockwise saccades (diff = -16.5°, CI = [-27.6, -8.8]). In each of the separate maps, the same general pattern is found in long-term memory. It is particularly pronounced in V1 (diff = -13.3, CI [-49.8, 9.9]), V3 (diff = -14.3°, CI [-17.5, -10.6]), V3ab (diff = -12.7°, CI [-20.4, -7.3]), LO1 (diff = -38.7°, CI [-69.1, -8]), and IPS0 (diff = -27.6°, CI [-43, -12.3]). In no map does the tuning go in the opposite direction of the saccades.

**Figure 7.**
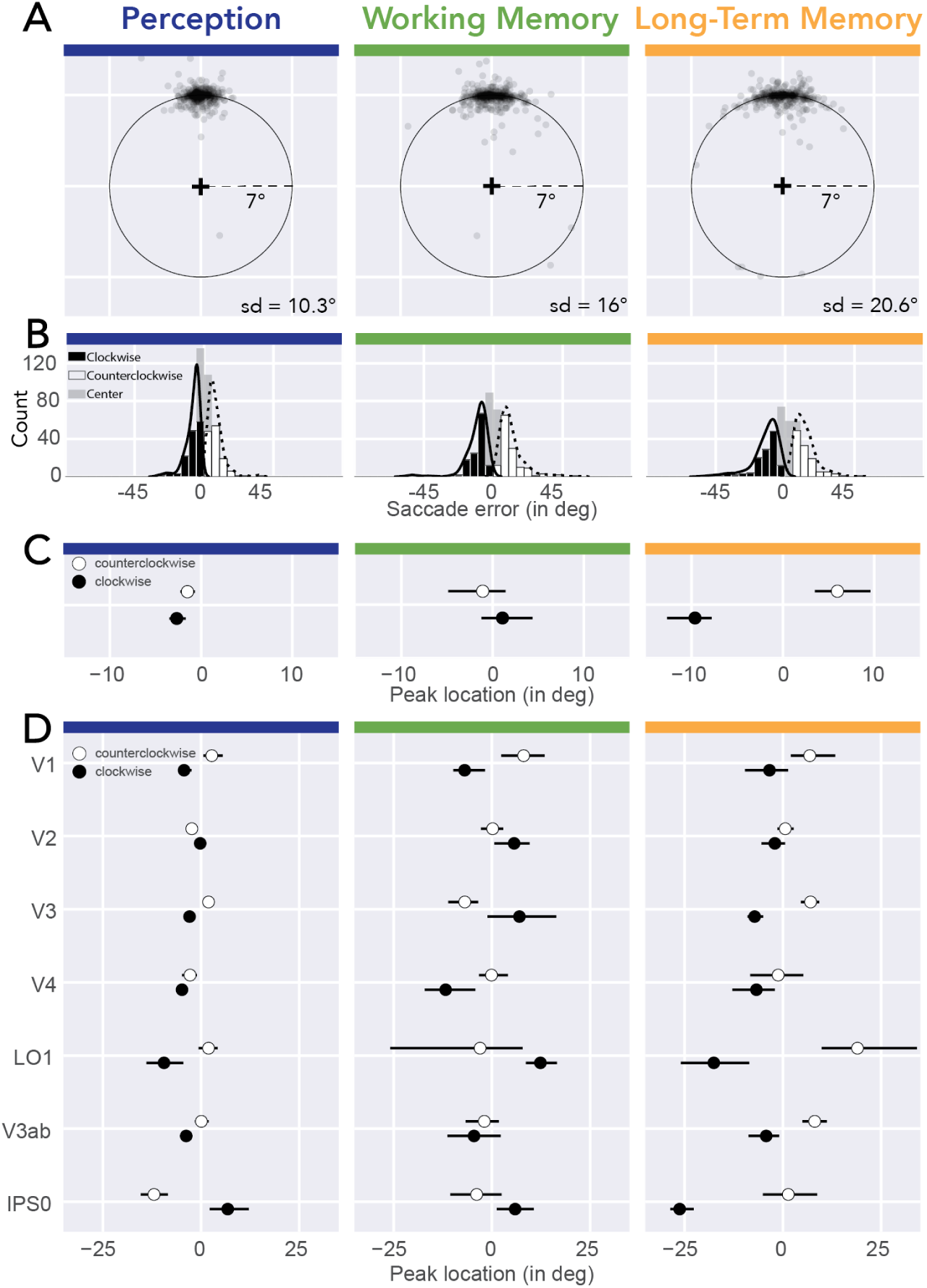
Saccade errors align with cortical tuning in long-term memory. A) Saccade response spread in visual space. We aligned saccade responses to the target location by subtracting their polar angle distance from target, and added 90° to align to the vertical axis. Saccade responses during memory had greater error along polar angle than during perception, consistent with the broader spatial tuning during memory within early visual maps. Standard deviation of saccade errors in polar angle are shown in the bottom right. B) Histograms of saccade error along polar angle. We grouped trials into tertiles: clockwise (black), center (gray), or counterclockwise (white) to target polar angle location. The tertiles were defined for each subject and condition, hence there is some overlap in the group-average histograms plotted. C) Peak location estimates for clockwise/counterclockwise trial groups. We fit polar angle activation profiles to the clockwise (black circles) and counterclockwise (white circles) groups. Error bars represent the bootstrapped 68% confidence intervals. D) Peak location estimates for clockwise/counterclockwise groups for each measured visual map. Error bars represent 68% confidence intervals. The panels here are generated using *fig6_09-16-2024.py*.

In contrast, for working memory, there was no systematic relationship between the peak estimates of spatial tuning and saccade direction for all rois as a group (diff = 3.1°, CI [-3.8, 12.2]), but there were decent effects in V1 (diff -14.4, CI [-27.5, -2.7]). And there was a small overall effect in perception (diff = -1.1°, CI [-2.5, 0.4]), particularly in V1 (diff = -7.2, CI [-13.9, -1.3]), V3 (diff = -4.7, CI [-7.2, -1.7]), V3ab (diff = -3.8, CI [-7.3, 0.5]), and LO1 (diff = -10.7, CI [-15.2, -4.3]). This could be due to a restricted range of saccade errors in perception, and to the possibility that when there are errors in working memory, they develop gradually over the trial, as the representation drifts away from the viewed target. Since the analysis of peak location pools across the whole delay, this would diminish the impact of biases near the ends of the trials.

The long-term memory results suggest that information reconstructed in visual maps is likely used for memory-guided behavior.

## Discussion

We measured spatial tuning of cortical responses during perception, working memory, and long-term memory. We found that both long-term and working memory scale the spatial tuning of responses in early but not late visual field maps. We further demonstrated the importance of memory-driven visual cortex activations by showing that they correlate with behavior: trial-to-trial variation in the spatial tuning of cortical responses during long-term memory is related to the accuracy of subsequent oculomotor responses.

### Spatial tuning in visual cortex during long-term memory differs from perception

Long-term memory retrieval of information from hippocampus can reinstate patterns of cortical activity evoked during encoding (Johnson et al., 2009; Pearson et al., 2015; Rugg et al., 2008; Wheeler et al., 2000). This ‘cortical reinstatement’ is supported by neurally-inspired models of episodic memory (Damasio, 1989; McClelland et al., 1995; Rolls, 2000; Teyler & Rudy, 2007). These theories do not imply that memory and perceptual representations are identical. For example, memory responses are noisier than perceptual responses. One possibility is that neural responses during memory reinstatement are like those during perception, except for decreased signal-to-noise level (Johnson et al., 2009; Pearson et al., 2015; Rugg et al., 2008; Wheeler et al., 2000).

Replicating two recent studies (Breedlove et al., 2020; Favila et al., 2022), we find only partial support for such reinstatement. On the one hand, spatial tuning during long-term memory retrieval peaks at the expected locations in multiple visual field maps, as early as V1, demonstrating that memory reinstatement is stimulus specific. On the other hand, long-term memory responses in earlier visual maps (V1-V3) were more broadly tuned than perceptual responses, demonstrating that memory driven-activity is not simply a reinstatement of perceptual responses. A likely explanation lies in the architecture of visual cortex: during feedback, earlier maps inherit the lower spatial precision characteristic of later visual maps where neurons have large receptive fields. Prior work supports this reversal of cortical information flow as an explanation for top-down reinstatement during imagery (<u>Kosslyn 1994; Dentico et al. 2014;</u> <u>Pearson 2019; Dijkstra et al. 2020; Breedlove et al. 2020</u>). Lower precision in later stages is expected from a hierarchical encoding process with increasing levels of tolerance to size and position of stimuli (Ito et al., 1995; Kay, Winawer, et al., 2013); the greater size and position tolerance in later visual maps reflects a loss of position information likely not recoverable during retrieval. This information loss puts a limit on the precision of cortical reinstatement, independent of the lower amplitude signals during memory.

### How much information is lost in long-term memory?

In addition to architectural constraints, the precision of top-down generated signals may also depend on task demands. A comparison between our results and those of Favila et al. (2022) supports this idea. While both studies showed broader tuning for memory than perception in early visual maps, the decreased precision during memory was more pronounced in Favila’s study than this one (Figure 8a). This difference is unlikely due to measurement noise, or differences in eccentricity (4° vs 7°) or analysis. We infer this because the tuning width estimates of the two studies match during perception and, in later maps, during memory - the only differences between them being long-term memory in V1-V3. We speculate that the difference arises from encoding demands. The studies differed in the number of remembered targets and their spacing: four targets spaced by 90 deg in Favila et al vs 16 targets spaced on average 22.5 deg here. The narrow spacing in our study likely required more precise encoding, leading to narrow tuning in V1-V3 during memory. A post-hoc analysis supports this interpretation. Due to the random placement of stimulus locations within polar angle bins, the spacing between our stimuli varied (Figure 8c). For some stimuli, the spacing was just a few degrees. For others it was over 30 degrees. Although two stimuli were never simultaneously present, the training protocol required saccade responses to be closer to the correct target than to any other target. Hence nearby targets required greater precision. In V1-V3, tuning was narrower for targets with near neighbors than with far neighbors (Figure 8b). Tuning width for far-spaced targets is similar to Favila et al’s results. The implication is that increased competition during memory training invokes a greater demand for spatial precision, resulting in narrower tuning during recall. Further studies are needed to test whether this effect is driven by hippocampal pattern separation (Bakker et al., 2008; Yassa & Stark, 2011), or by selective attention, either at encoding or retrieval.

**Figure 8.**
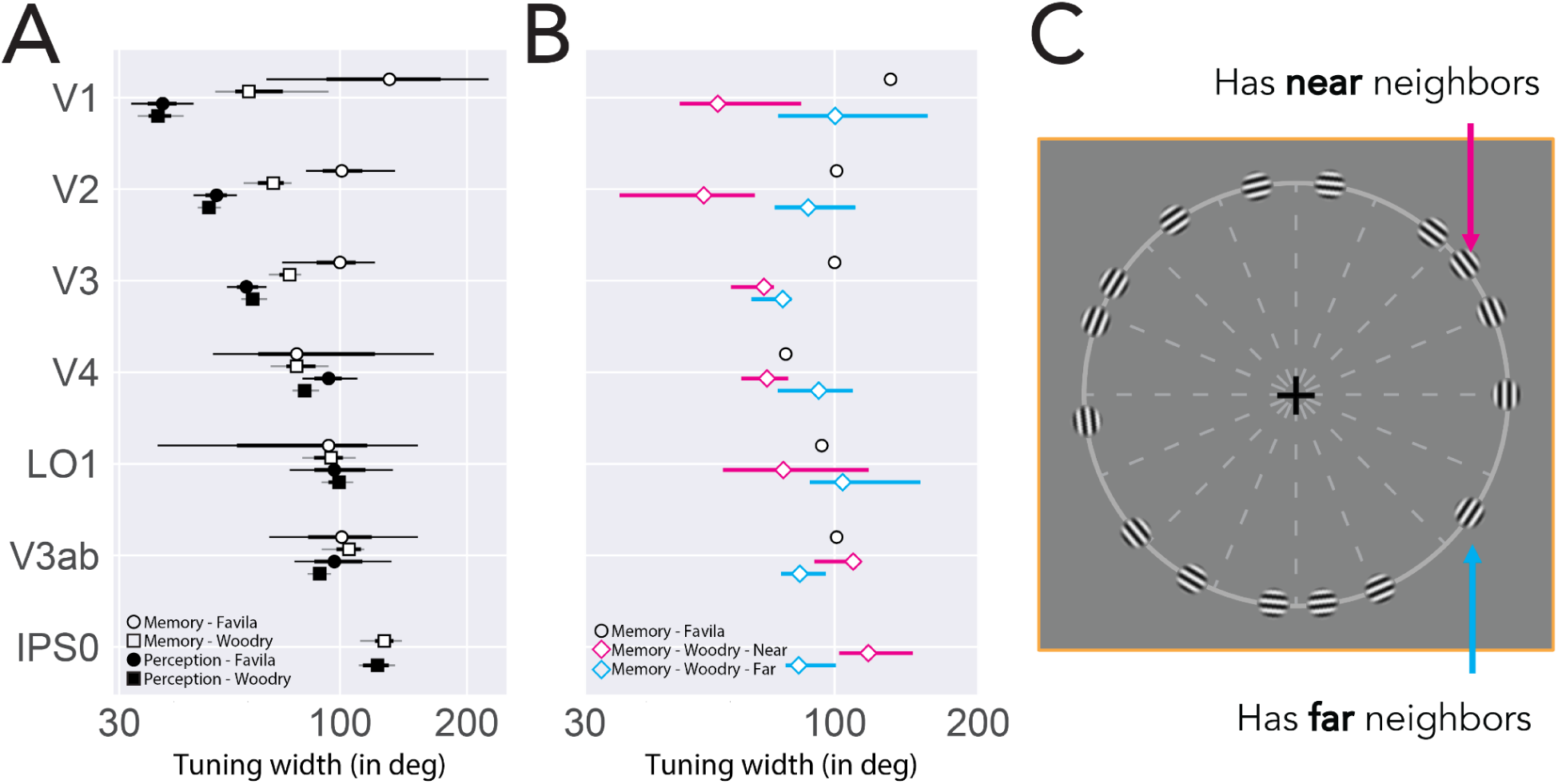
Systematic differences in spatial tuning of cortical responses. A) Comparison of spatial tuning (FWHM) estimates across measured visual maps between Favila et al. (2022; circles) and our study (squares). Both studies show large agreement in their spatial tuning estimates during the perception condition (filled), and in later visual maps V4-IPS0 during long-term memory retrieval (unfilled). During long-term memory, tuning widths differ in early visual maps V1-V3. The main differences between ours and the Favila study’s memory conditions are in the spacing of the stimuli; their study required the retrieval of four targets equally spaced around the visual field, whereas ours involved the retrieval of 16 targets that varied in their separation. B) Spatial tuning width estimates across visual maps, split by whether targets were spaced near to (pink diamonds; spacing: 1.7° - 12°, mean: 7.7°, std: 3.3°) or spaced far from (blue diamonds; spacing: 19.2° - 31.8°, mean: 23°, std: 3.4°) each other. Also plotted are the mean tuning width estimates from both the Favila et al. study (circles) and this study (squares). The tuning widths from far-spaced targets in our study more closely resemble the broader tuning observed in the Favila et al. study, suggesting the differences between mean width estimates between our studies is in large part due to the increased precision required of more closely spaced targets. C) Example of this study’s long-term memory retrieval targets. 16 Target stimuli, each sampled from within 22.5° bins, vary in their separation from other targets. Some targets were spaced far apart, while others were spaced near each other, meaning those spaced near to each other likely required more precision during later recall. This is reflected in the narrower spatial tuning observed during these trials than for the “far-spaced” trials. Panels A and B are generated using *fig7_09-16-2024.py*.

### Amplitude and tuning dynamics in visual cortex during memory

A peripheral visual stimulus occurred at the beginning of working memory but not long-term memory trials. During working memory, the location of this stimulus must be maintained over the delay period such that a memory-guided saccade can be generated to the stored location. During long-term memory, the target of the memory-guided saccade was retrieved from memory and not based on a visual stimulus. Consequently, in all areas a robust transient response time-locked to the early visual stimulus distinguished working and long-term memory (fig. 6b). In early areas, this transient response was narrowly tuned, matched to the perception condition. If working memory simply maintained recent activity, one would expect the tuning to stay narrow throughout the delay, but it does not. In fact, the tuning widths of responses in visual cortex during the later epochs of the trial were remarkably similar during working and long-term memory (fig. 6d). Despite originating from distinct bottom-up visual and top-down mnemonic sources, perhaps the contents of memory are similar by this time (i.e., the target of the memory-guided saccade). Indeed, visual memories flexibly and efficiently represent visual properties that are most behaviorally relevant (Chunharas et al., 2023; Duan & Curtis, 2024; Kwak & Curtis, 2022; Li & Curtis, 2023), in this case the goal of the memory-guided saccade. What limits the precision appears to be not whether the stimulus was recently viewed (working memory) or retrieved from long-term storage, but rather that the stimulus representation is maintained in the absence of perceptual input.

Surprisingly, responses in V1-V3 were more precise in some long-term memory trials (near spaced stimuli, see fig. 8b) than working memory, despite the longer delay between encoding and retrieval for long-term memory. This is likely because during our long-term memory experiment, subjects viewed the stimuli many times (about 60 times each during pre-scan training), and for near-spaced stimuli, needed to make fine distinctions. Recent work shows that when stimuli are highly learned in long-term memory, the working memory of these stimuli becomes more precise (Miller et al., 2022), similar to the relatively precise representations in V1-V3 for our near spaced long-term memory stimuli.

### Are the memory responses in visual cortex behaviorally relevant?

Our trial-by-trial analyses demonstrated an alignment between behavioral responses and cortical activation in long-term memory, but not during perception or working memory. Other studies have reported links between trial-to-trial variability in long-term memory responses and self-reported vividness (Bone et al., 2019) and recognition accuracy (Bone et al., 2020). These studies do not predict the specific behavioral errors based on the cortical responses. Quantifying neural responses in stimulus units (polar angle) enables us to tightly link the trial-by-trial responses to behavior: the direction of the error in cortical responses is predictive of the direction of the error in behavior.

Nonetheless, our results are correlative. Demonstrating the *necessity* of visual cortex activity for behavior requires manipulating the neural responses. A recent TMS study suggests that disrupting V1 activity during spatial working memory maintenance reduces memory-guided saccade accuracy (Dake & Curtis, 2024), though effects of TMS on V1 during WM delays did not reduce accuracy for feature judgments (orientation - <u>Rademaker et al. 2017</u>; color - <u>van</u> <u>Lamsweerde and Johnson 2017</u>). Perhaps it is surprising then that we only find a correspondence between cortical activity and behavior in long-term memory, not working memory. It is unclear why there is a difference between the conditions. One likely possibility is that the saccade errors in working memory are smaller, making it more difficult to separate encoding errors from motor noise. Overall, the combination of a correspondence between behavior and long-term memory cortical responses reported here, between behavior and working memory cortical responses reported elsewhere <u>(Ester et al. 2013; van Bergen et al. 2015; Hallenbeck et al. 2021; Li and</u> <u>Curtis 2023)</u>, and between TMS and behavior (Dake & Curtis, 2024) make a strong case that memory responses in early visual cortex are relevant for behavior.

### Is it just attention?

Memory shares features with attention, in that both involve modulation of neural responses that are not purely stimulus-driven. There are shared mechanisms between attention and long-term memory (Aly & Turk-Browne, 2017) and attention and working memory (Jerde et al., 2012; Panichello & Buschman, 2021; Zhou et al., 2022). Indeed, one way to describe sustaining a memory, whether sustained from recent exposure or retrieved from a long-term store, is attending to an internal feature (Cabeza et al., 2008). Thus our memory tasks surely entail some elements of attention. Nonetheless, there are no results from the attention literature that would have enabled us to predict the pattern of tuning widths we observe in early vs late visual areas among perception, working memory, and long-term memory. Moreover, attention alone could not produce cortical patterns of activation in visual cortex spatially tuned to the location of a remembered target. In the absence of any visual stimulation at that location during the long-term memory trials, the location information must first be retrieved from memory storage. Further experiments are needed to disentangle the contributions of attention and episodic memory to activation patterns in visual cortex.

### Decoding vs encoding approaches to studying memory representations

Our use of retinotopic models allowed us to quantify and compare sensory activation during perception, working memory, and long-term memory in terms of spatial tuning functions. This differs from the decoding approach, often used for stimulus orientation or other stimulus features which do not vary systematically at the mm to cm scale of fMRI measurement. Decoding fMRI responses in memory or imagery is less accurate than perception (Naselaris et al., 2015). Such results have been coupled to the idea that memory or imagery representations are like weak forms of perception, similar to viewing stimuli at low contrast (Pearson, 2019). But the decrease in decoding accuracy can arise from a decrease in signal-to-noise ratio (expected from a weaker stimulus) or a decrease in precision, or a combination. Our stimulus manipulation allowed us to quantify precision independent of amplitude. Precision was lower in memory, and also varied with task demands and map location in the visual hierarchy. Separating precision of the representation from the signal-to-noise ratio is difficult for stimulus features whose representations vary over a sub-mm scale, such as orientation, despite recent advances in studying their encoding (Gardner & Liu, 2019; Roth et al., 2018; Sprague et al., 2018). In contrast, it is straightforward in the spatial domain, making spatial manipulations a useful tool for comparing neural representations across conditions. But spatial manipulations are not just a tool. Space is a fundamental part of visual representations (Wandell & Winawer, 2011) and its role in both perception and memory disorders remains a central topic in cognitive neuroscience (Bisiach & Luzzatti, 1978; Farah, 2003).

## Acknowledgements

We thank New York University’s Center for Brain Imaging for technical support. This research was supported by the National Eye Institute (R01 EY033925 & R01 EY016407 to C.E. Curtis; R01 EY027401 to J. Winawer; P30 EY013079 Core grant for vision) by the National Institute of Mental Health (R01 MH111417 to J. Winawer), and pilot funds from the NYU Center for Brain Imaging to R. Woodry. This work was supported in part through the NYU IT High Performance Computing resources, services, and staff expertise. We thank Ilona Bloem, Jan Kurzawski, Marc Himmelberg, Rania Ezzo, and Ekin Tunçok for help with scanning; Serra Favila, Tommy Sprague, and Ekin Tunçok for helpful comments on the manuscript; and Serra Favila for sharing data and code. Because this is my first submission I‘d like to extend special thanks to Kate Yurgil and Liz Chrastil for taking a chance on me.

## References

Abraham, A., Pedregosa, F., Eickenberg, M., Gervais, P., Mueller, A., Kossaifi, J., Gramfort, A., Thirion, B., & Varoquaux, G. (2014). Machine learning for neuroimaging with scikit-learn. Frontiers in Neuroinformatics, 8, 14.

Albers, A. M., Kok, P., Toni, I., Dijkerman, H. C., & de Lange, F. P. (2013). Shared representations for working memory and mental imagery in early visual cortex. Current Biology: CB, 23(15), 1427–1431.

Allen, E. J., St-Yves, G., Wu, Y., Breedlove, J. L., Prince, J. S., Dowdle, L. T., Nau, M., Caron, B., Pestilli, F., Charest, I., Hutchinson, J. B., Naselaris, T., & Kay, K. (2021). A massive 7T fMRI dataset to bridge cognitive neuroscience and artificial intelligence. Nature Neuroscience, 25(1), 116–126.

Aly, M., & Turk-Browne, N. B. (2017). How hippocampal memory shapes, and is shaped by, attention. In D. E. Hannula (Ed.), The hippocampus from cells to systems: Structure, connectivity, and functional contributions to memory and flexible cognition (pp (Vol. 589, pp. 369–403).

Avants, B. B., Epstein, C. L., Grossman, M., & Gee, J. C. (2008). Symmetric diffeomorphic image registration with cross-correlation: evaluating automated labeling of elderly and neurodegenerative brain. Medical Image Analysis, 12(1), 26–41.

Bakker, A., Kirwan, C. B., Miller, M., & Stark, C. E. L. (2008). Pattern separation in the human hippocampal CA3 and dentate gyrus. Science, 319(5870), 1640–1642.

Benson, N. C., & Winawer, J. (2018). Bayesian analysis of retinotopic maps. eLife, 7. 10.7554/eLife.40224

Benson, N. C., Yoon, J. M. D., Forenzo, D., Engel, S. A., Kay, K. N., & Winawer, J. (2022). Variability of the Surface Area of the V1, V2, and V3 Maps in a Large Sample of Human Observers. The Journal of Neuroscience: The Official Journal of the Society for Neuroscience, 42(46), 8629–8646.

Bisiach, E., & Luzzatti, C. (1978). Unilateral neglect of representational space. Cortex; a Journal Devoted to the Study of the Nervous System and Behavior, 14(1), 129–133.

Bone, M. B., Ahmad, F., & Buchsbaum, B. R. (2020). Feature-specific neural reactivation during episodic memory. Nature Communications, 11(1), 1945.

Bone, M. B., St-Laurent, M., Dang, C., McQuiggan, D. A., Ryan, J. D., & Buchsbaum, B. R. (2019). Eye Movement Reinstatement and Neural Reactivation During Mental Imagery. Cerebral Cortex, 29(3), 1075–1089.

Bosch, S. E., Jehee, J. F. M., Fernández, G., & Doeller, C. F. (2014). Reinstatement of associative memories in early visual cortex is signaled by the hippocampus. The Journal of Neuroscience: The Official Journal of the Society for Neuroscience, 34(22), 7493–7500.

Breedlove, J. L., St-Yves, G., Olman, C. A., & Naselaris, T. (2020). Generative Feedback Explains Distinct Brain Activity Codes for Seen and Mental Images. Current Biology: CB, 30(12), 2211–2224.e6.

Brett, M., Markiewicz, C. J., Hanke, M., Côté, M.-A., Cipollini, B., McCarthy, P., Jarecka, D., Cheng, C. P., Halchenko, Y. O., Cottaar, M., Larson, E., Ghosh, S., Wassermann, D., Gerhard, S., Lee, G. R., Wang, H.-T., Kastman, E., Kaczmarzyk, J., Guidotti, R., … freec84. (2022). nipy/nibabel: 3.2.2 (3.2.2). Zenodo. 10.5281/zenodo.6617121

Brodeur, M. B., Dionne-Dostie, E., Montreuil, T., & Lepage, M. (2010). The Bank of Standardized Stimuli (BOSS), a new set of 480 normative photos of objects to be used as visual stimuli in cognitive research. PloS One, 5(5), e10773.

Cabeza, R., Ciaramelli, E., Olson, I. R., & Moscovitch, M. (2008). The parietal cortex and episodic memory: an attentional account. Nature Reviews. Neuroscience, 9(8), 613–625.

Chunharas, C., Hettwer, M. D., Wolff, M. J., & Rademaker, R. L. (2023). A gradual transition from veridical to categorical representations along the visual hierarchy during working memory, but not perception. bioRxiv.org: The Preprint Server for Biology. 10.1101/2023.05.18.541327

Cox, R. W., & Hyde, J. S. (1997). Software tools for analysis and visualization of fMRI data. NMR in Biomedicine, 10(4-5), 171–178.

Curtis, C. E., & D’Esposito, M. (2003). Persistent activity in the prefrontal cortex during working memory. Trends in Cognitive Sciences, 7(9), 415–423.

Curtis, C. E., & Sprague, T. C. (2021). Persistent Activity During Working Memory From Front to Back. Frontiers in Neural Circuits, 15, 696060.

Dake, M., & Curtis, C. E. (2024). Perturbing human V1 degrades the precision of visual working memory. Biorxiv.

Dale, A. M., Fischl, B., & Sereno, M. I. (1999). Cortical surface-based analysis. I. Segmentation and surface reconstruction. NeuroImage, 9(2), 179–194.

Damasio, A. R. (1989). Time-locked multiregional retroactivation: a systems-level proposal for the neural substrates of recall and recognition. Cognition, 33(1-2), 25–62.

D’Esposito, M., & Postle, B. R. (2015). The cognitive neuroscience of working memory. Annual Review of Psychology, 66, 115–142.

Duan, Z., & Curtis, C. E. (2024). Visual working memories are abstractions of percepts. eLife, 13. 10.7554/eLife.94191

Dumoulin, S. O., & Wandell, B. A. (2008). Population receptive field estimates in human visual cortex. NeuroImage, 39(2), 647–660.

Esteban, O., Blair, R., Markiewicz, C. J., Berleant, S. L., Moodie, C., Ma, F., Isik, A. I., Erramuzpe, A., Kent, J. D., Goncalves, M., & Others. (2018). fMRIPrep. Software. Zenodo.

Esteban, O., Markiewicz, C. J., Blair, R. W., Moodie, C. A., Isik, A. I., Erramuzpe, A., Kent, J. D., Goncalves, M., DuPre, E., Snyder, M., Oya, H., Ghosh, S. S., Wright, J., Durnez, J., Poldrack, R. A., & Gorgolewski, K. J. (2019). fMRIPrep: a robust preprocessing pipeline for functional MRI. Nature Methods, 16(1), 111–116.

Farah, M. J. (2003). Disorders of visual-spatial perception and cognition. Clinical Neuropsychology, 4. https://books.google.com/books?hl=en&lr=&id=MT_RCwAAQBAJ&oi=fnd&pg=PA152&dq=Farah+and+Epstein+2003&ots=-pQhhikHZo&sig=uSlYDe0ibIeT4wfc7toVl8DFHmM

Favila, S. E., Kuhl, B. A., & Winawer, J. (2022). Perception and memory have distinct spatial tuning properties in human visual cortex. Nature Communications, 13(1), 5864.

Gardner, J. L., & Liu, T. (2019). Inverted Encoding Models Reconstruct an Arbitrary Model Response, Not the Stimulus. eNeuro, 6(2). 10.1523/ENEURO.0363-18.2019

Gorgolewski, K., Burns, C. D., Madison, C., Clark, D., Halchenko, Y. O., Waskom, M. L., & Ghosh, S. S. (2011). Nipype: a flexible, lightweight and extensible neuroimaging data processing framework in python. Frontiers in Neuroinformatics, 5, 13.

Gorgolewski, K. J., Auer, T., Calhoun, V. D., Craddock, R. C., Das, S., Duff, E. P., Flandin, G., Ghosh, S. S., Glatard, T., Halchenko, Y. O., Handwerker, D. A., Hanke, M., Keator, D., Li, X., Michael, Z., Maumet, C., Nichols, B. N., Nichols, T. E., Pellman, J., … Poldrack, R. A. (2016). The brain imaging data structure, a format for organizing and describing outputs of neuroimaging experiments. Scientific Data, 3(1), 1–9.

Gorgolewski, K. J., Nichols, T., Kennedy, D. N., Poline, J.-B., & Poldrack, R. A. (2018). Making replication prestigious. The Behavioral and Brain Sciences, 41, e131.

Greve, D. N., & Fischl, B. (2009). Accurate and robust brain image alignment using boundary-based registration. NeuroImage, 48(1), 63–72.

Hallenbeck, G. E., Sprague, T. C., Rahmati, M., Sreenivasan, K. K., & Curtis, C. E. (2021). Working memory representations in visual cortex mediate distraction effects. Nature Communications, 12(1), 4714.

Halchenko, Y. O., et al. Open Source Software: Heudiconv. Zenodo, 10.5281/zenodo.1306159 (2018).

Harrison, S. A., & Tong, F. (2009). Decoding reveals the contents of visual working memory in early visual areas. Nature, 458(7238), 632–635.

Himmelberg, M. M., Kurzawski, J. W., Benson, N. C., Pelli, D. G., Carrasco, M., & Winawer, J. (2021). Cross-dataset reproducibility of human retinotopic maps. NeuroImage, 244, 118609.

Hunter J, “Matplotlib: A 2D Graphics Environment”, Computing in Science & Engineering, vol. 9, no. 3, pp. 90–95, 2007

Inouye, T. (1909). Die Sehstörungen bei Schussverletzungen der kortikalen Sehsphäre: nach Beobachtungen an Verwundeten der letzten japanischen Kriege. Engelmann.

Ishai, A., & Sagi, D. (1995). Common mechanisms of visual imagery and perception. Science, 268(5218), 1772–1774.

Ishai, A., & Sagi, D. (1997). Visual imagery facilitates visual perception: psychophysical evidence. Journal of Cognitive Neuroscience, 9(4), 476–489.

Ito, M., Tamura, H., Fujita, I., & Tanaka, K. (1995). Size and position invariance of neuronal responses in monkey inferotemporal cortex. Journal of Neurophysiology, 73(1), 218–226.

Jenkinson, M., Bannister, P., Brady, M., & Smith, S. (2002). Improved optimization for the robust and accurate linear registration and motion correction of brain images. NeuroImage, 17(2), 825–841.

Jerde, T. A., Merriam, E. P., Riggall, A. C., Hedges, J. H., & Curtis, C. E. (2012). Prioritized maps of space in human frontoparietal cortex. The Journal of Neuroscience: The Official Journal of the Society for Neuroscience, 32(48), 17382–17390.

Johnson, J. D., McDuff, S. G. R., Rugg, M. D., & Norman, K. A. (2009). Recollection, familiarity, and cortical reinstatement: a multivoxel pattern analysis. Neuron, 63(5), 697–708.

Kay, K. N., Rokem, A., Winawer, J., Dougherty, R. F., & Wandell, B. A. (2013). GLMdenoise: a fast, automated technique for denoising task-based fMRI data. Frontiers in Neuroscience, 7, 247.

Kay, K. N., Winawer, J., Mezer, A., & Wandell, B. A. (2013). Compressive spatial summation in human visual cortex. Journal of Neurophysiology, 110(2), 481–494.

Klein, A. (2017). Mindboggle-101 templates (unlabeled images from a population of brains). Harvard Dataverse.

Kosslyn, S. M., Thompson, W. L., Kim, I. J., & Alpert, N. M. (1995). Topographical representations of mental images in primary visual cortex. Nature, 378(6556), 496–498.

Kwak, Y., & Curtis, C. E. (2022). Unveiling the abstract format of mnemonic representations. Neuron, 110(11), 1822–1828.e5.

Lanczos, C. (1964). Evaluation of Noisy Data. Journal of the Society for Industrial and Applied Mathematics Series B Numerical Analysis, 1(1), 76–85.

Larsson, J., & Heeger, D. J. (2006). Two retinotopic visual areas in human lateral occipital cortex. The Journal of Neuroscience: The Official Journal of the Society for Neuroscience, 26(51), 13128–13142.

Li, H.-H., & Curtis, C. E. (2023). Neural population dynamics of human working memory. Current Biology: CB, 33(17), 3775–3784.e4.

Li, H.-H., Sprague, T. C., Yoo, A. H., Ma, W. J., & Curtis, C. E. (2021). Joint representation of working memory and uncertainty in human cortex. Neuron, 109(22), 3699–3712.e6.

Mackey, W. E., Winawer, J., & Curtis, C. E. (2017). Visual field map clusters in human frontoparietal cortex. eLife, 6. 10.7554/eLife.22974

McClelland, J. L., McNaughton, B. L., & O’Reilly, R. C. (1995). Why there are complementary learning systems in the hippocampus and neocortex: insights from the successes and failures of connectionist models of learning and memory. Psychological Review, 102(3), 419–457.

Miller, J. A., Tambini, A., Kiyonaga, A., & D’Esposito, M. (2022). Long-term learning transforms prefrontal cortex representations during working memory. Neuron, 110(22), 3805–3819.e6.

Naselaris, T., Olman, C. A., Stansbury, D. E., Ugurbil, K., & Gallant, J. L. (2015). A voxel-wise encoding model for early visual areas decodes mental images of remembered scenes. NeuroImage, 105, 215–228.

The pandas development team. (2024). pandas-dev/pandas: Pandas (v2.2.1). Zenodo. 10.5281/zenodo.10697587

Panichello, M. F., & Buschman, T. J. (2021). Shared mechanisms underlie the control of working memory and attention. Nature, 592(7855), 601–605.

Pearson, J. (2019). The human imagination: the cognitive neuroscience of visual mental imagery. Nature Reviews. Neuroscience, 20(10), 624–634.

Pearson, J., Naselaris, T., Holmes, E. A., & Kosslyn, S. M. (2015). Mental Imagery: Functional Mechanisms and Clinical Applications. Trends in Cognitive Sciences, 19(10), 590–602.

Pedregosa, F., Varoquaux, G., Gramfort, A., Michel, V., Thirion, B., Grisel, O., Blondel, M., Louppe, G., Prettenhofer, P., Weiss, R., Weiss, R. J., Vanderplas, J., Passos, A., Cournapeau, D., Brucher, M., Perrot, M., & Duchesnay, E. (2011). Scikit-learn: Machine Learning in Python. Journal of Machine Learning Research: JMLR, abs/1201.0490. 10.5555/1953048.2078195

Prince, J. S., Charest, I., Kurzawski, J. W., Pyles, J. A., Tarr, M. J., & Kay, K. N. (2022). Improving the accuracy of single-trial fMRI response estimates using GLMsingle. eLife, 11. 10.7554/eLife.77599

Rademaker, R. L., Chunharas, C., & Serences, J. T. (2019). Coexisting representations of sensory and mnemonic information in human visual cortex. Nature Neuroscience, 22(8), 1336–1344.

Reuter, M., Rosas, H. D., & Fischl, B. (2010). Highly accurate inverse consistent registration: a robust approach. NeuroImage, 53(4), 1181–1196.

Rolls, E. T. (2000). Hippocampo-cortical and cortico-cortical backprojections. Hippocampus, 10(4), 380–388.

Roth, Z. N., Heeger, D. J., & Merriam, E. P. (2018). Stimulus vignetting and orientation selectivity in human visual cortex. eLife, 7. 10.7554/eLife.37241

Rugg, M. D., Johnson, J. D., Park, H., & Uncapher, M. R. (2008). Encoding-retrieval overlap in human episodic memory: a functional neuroimaging perspective. Progress in Brain Research, 169, 339–352.

Schacter, D. L., Norman, K. A., & Koutstaal, W. (1998). The cognitive neuroscience of constructive memory. Annual Review of Psychology, 49, 289–318.

Serences, J. T. (2016). Neural mechanisms of information storage in visual short-term memory. In Vision Research (Vol. 128, pp. 53–67). 10.1016/j.visres.2016.09.010

Serences, J. T., Ester, E. F., Vogel, E. K., & Awh, E. (2009). Stimulus-specific delay activity in human primary visual cortex. Psychological Science, 20(2), 207–214.

Smith, A. T., Singh, K. D., Williams, A. L., & Greenlee, M. W. (2001). Estimating receptive field size from fMRI data in human striate and extrastriate visual cortex. Cerebral Cortex, 11(12), 1182–1190.

Sprague, T. C., Adam, K. C. S., Foster, J. J., Rahmati, M., Sutterer, D. W., & Vo, V. A. (2018). Inverted Encoding Models Assay Population-Level Stimulus Representations, Not Single-Unit Neural Tuning. eNeuro, 5(3). 10.1523/ENEURO.0098-18.2018

Sprague, T. C., & Serences, J. T. (2013). Attention modulates spatial priority maps in the human occipital, parietal and frontal cortices. Nature Neuroscience, 16(12), 1879–1887.

Teyler, T. J., & Rudy, J. W. (2007). The hippocampal indexing theory and episodic memory: updating the index. Hippocampus, 17(12), 1158–1169.

Tong, F., Nakayama, K., Vaughan, J. T., & Kanwisher, N. (1998). Binocular rivalry and visual awareness in human extrastriate cortex. Neuron, 21(4), 753–759.

Tulving, E., & Thomson, D. M. (1973). Encoding specificity and retrieval processes in episodic memory. Psychological Review, 80(5), 352–373.

Tustison, N. J., Avants, B. B., Cook, P. A., Zheng, Y., Egan, A., Yushkevich, P. A., & Gee, J. C. (2010). N4ITK: improved N3 bias correction. IEEE Transactions on Medical Imaging, 29(6), 1310–1320.

van Bergen, R. S., Ma, W. J., Pratte, M. S., & Jehee, J. F. M. (2015). Sensory uncertainty decoded from visual cortex predicts behavior. Nature Neuroscience, 18(12), 1728–1730.

Van Essen, D. C., & Maunsell, J. H. R. (1983). Hierarchical organization and functional streams in the visual cortex. Trends in Neurosciences, 6, 370–375.

Virtanen, P., Gommers, R., Oliphant, T. E., Haberland, M., Reddy, T., Cournapeau, D., Burovski, E., Peterson, P., Weckesser, W., Bright, J., van der Walt, S. J., Brett, M., Wilson, J., Millman, K. J., Mayorov, N., Nelson, A. R. J., Jones, E., Kern, R., Larson, E., … van Mulbregt, P. (2020). SciPy 1.0: fundamental algorithms for scientific computing in Python. Nature Methods, 17(3), 261–272.

Vo, V. A., Sutterer, D. W., Foster, J. J., Sprague, T. C., Awh, E., & Serences, J. T. (2022). Shared Representational Formats for Information Maintained in Working Memory and Information Retrieved from Long-Term Memory. Cerebral Cortex, 32(5), 1077–1092.

Wandell, B. A., & Winawer, J. (2011). Imaging retinotopic maps in the human brain. Vision Research, 51(7), 718–737.

Waskom, M. L., (2021). seaborn: statistical data visualization. Journal of Open Source Software, 6(60), 3021, 10.21105/joss.03021

Wheeler, M. E., Petersen, S. E., & Buckner, R. L. (2000). Memory’s echo: vivid remembering reactivates sensory-specific cortex. Proceedings of the National Academy of Sciences of the United States of America, 97(20), 11125–11129.

Winawer, J., & Witthoft, N. (2015). Human V4 and ventral occipital retinotopic maps. Visual Neuroscience, 32, E020.

Xu, Y. (2017). Reevaluating the Sensory Account of Visual Working Memory Storage. Trends in Cognitive Sciences, 21(10), 794–815.

Yassa, M. A., & Stark, C. E. L. (2011). Pattern separation in the hippocampus. Trends in Neurosciences, 34(10), 515–525.

Zhang, Y., Brady, J. M., & Smith, S. M. (2001). An hmrf-em algorithm for partial volume segmentation of brain mri fmrib technical report tr01yz1. Brain: A Journal of Neurology. https://www.fmrib.ox.ac.uk/datasets/techrep/tr01yz1/tr01yz1.pdf

Zhou, Y., Curtis, C. E., Sreenivasan, K. K., & Fougnie, D. (2022). Common Neural Mechanisms Control Attention and Working Memory. The Journal of Neuroscience: The Official Journal of the Society for Neuroscience, 42(37), 7110–7120.

